# The CHCHD2-CHCHD10 protein complex is modulated by mitochondrial dysfunction and alters lipid homeostasis in the mouse brain

**DOI:** 10.1101/2024.09.10.612325

**Authors:** Jule Gerlach, Paola Pireddu, Xiaoqun Zhang, Simon Wetzel, Mara Mennuni, Dusanka Milenkovic, Hendrik Nolte, Niclas Branzell, Ibrahim Kaya, Rodolfo Garcia Villegas, Diana Rubalcava-Gracia, David Alsina, Regina Feederle, Per E. Andrén, Thomas Langer, Per Svenningsson, Roberta Filograna

## Abstract

The highly conserved CHCHD2 and CHCHD10 are small mitochondrial proteins residing in the intermembrane space. Recently, mutations in the *CHCHD2* and *CHCHD10* genes have been linked to severe disorders, including Parkinson’s disease and amyotrophic lateral sclerosis. In cultured cells, a small fraction of CHCHD2 and CHCHD10 oligomerize to form a high molecular weight complex of unknown function. Here, we generated a whole-body *Chchd2* knockout mouse to investigate the *in vivo* role of CHCHD2 and its protein complex. We show that CHCHD2 is crucial for sustaining full motor capacity, normal striatal dopamine levels, and lipid homeostasis in the brain of adult male mice. We also demonstrate that in mouse tissues, CHCHD2 and CHCHD10 exist exclusively as a high molecular weight complex, whose levels are finely tuned under physiological conditions. In response to mitochondrial dysfunction, the abundance and size of the CHCHD2-CHCHD10 complex increases, a mechanism conserved across different tissues. Although the loss of CHCHD2 does not abolish CHCHD10 oligomerization, it enhances cell vulnerability to mitochondrial stress, suggesting that CHCHD2 is protective against mitochondrial damage. Our findings uncover the role of CHCHD2 in preserving tissue homeostasis and provide important insights into the involvement of the CHCHD2-CHCHD10 complex in human diseases.

## Introduction

Mitochondria arose from the engulfment of free-living α-proteobacterium by an archaeal host cell and evolved into essential organelles responsible for energy conversion and numerous other cellular functions. Mitochondria have a complex architecture with specialized compartments: the outer mitochondrial membrane (OMM) serves as the boundary to the cytosol, while the inner mitochondrial membrane (IMM), with its extensive folding and invaginations known as cristae, encloses the matrix and creates the intermembrane space (IMS). The IMS is a small aqueous compartment housing ~ 10% of the mammalian mitochondrial proteome ^1^, which coordinates several mitochondrial activities and whose functions rarely correspond to processes occurring in prokaryotes ^2^. In the past decade, it has become clear that the proteins residing in the IMS have diverse functions, including the exchange of proteins, lipids, or metal ions between the matrix and the cytosol, the initiation of apoptotic cascades, and the assembly of some the respiratory chain complexes ^3,4^.

All IMS proteins are encoded by the nuclear genome, synthesized in the cytosol and imported into mitochondria. The identification of the mitochondrial import machineries has led to the discovery of proteins that are imported into the IMS via the MIA pathway, which catalyzes protein folding through a disulfide relay system ^5^. Classical Mia40 substrates are evolutionarily conserved small mitochondrial proteins (usually <25 kDa) characterized by twin cysteine motifs. Some of these proteins belong to the coiled-coil-helix-coiled-coil-helix (CHCH) domain family and are emerging as important regulators of OXPHOS, lipid homeostasis, mitochondrial ultrastructure and dynamics ^6^. For example, CHCHD3 and CHCHD6, known as MIC19 and MIC25, are key components of the mitochondrial contact site and cristae organizing system (MICOS) complex ^7^. Mutations affecting proteins carrying the CHCH-domain have been associated with a broad spectrum of human diseases including cancer, obesity, myopathy and neurodegenerative disorders ^6^.

In 2015, heterozygous mutations in the *CHCHD2* gene were identified in three Japanese families with late-onset Parkinson’s disease (PD) ^8^. Further genetic studies discovered additional mutations linked to late-onset PD, juvenile PD, dementia with Lewy bodies and sporadic PD ^9–12^. Although rare, more than twenty disease-associated *CHCHD2* variants have been described to date (see review^13^). Most of these mutations are autosomal-dominant, indicating a gain-of-function mechanism of toxicity due to the accumulation of misfolded proteins ^14^. However, the discovery of nonsense and recessive *CHCHD2* mutations, causing protein truncation or decreased steady-state levels ^12,15,16^, supports a pathological mechanism driven by haploinsufficiency.

In humans, CHCHD2 is a 151 amino acid protein presenting 58% sequence identity with the family member CHCHD10 ^17^. Studies in human fibroblasts and HEK293T cells have provided solid evidence of a physical interaction between CHCHD2 and CHCHD10 in the IMS ^18,19^, where a fraction of these proteins oligomerize and assemble into a large protein complex ^18^. Notably, mutations in *CHCHD10* have been identified in patients with amyotrophic lateral sclerosis (ALS) and frontotemporal dementia ^20,21^, mitochondrial myopathy ^22^, spinal muscular atrophy ^23^, and Charcot-Marie-Tooth disease ^24^.

Extensive research conducted using *in vivo* and *in vitro* models has expanded our understanding of CHCHD2 and CHCHD10 but has also produced conflicting outcomes. Initial reports described CHCHD2 and CHCHD10 as MICOS subunits ^25,26^, while more recent data have largely disproved this hypothesis ^18^ arguing that CHCHD2 and CHCHD10 mutants may affect mitochondrial cristae architecture and only indirectly compromise MICOS stability ^27^. The loss of the CHCHD2/CHCHD10 ortholog in flies leads to a reduction in mitochondrial oxygen consumption and alterations in cristae structure ^28^. In contrast, CHCHD2 ablation in mice did not produce consistent phenotypes. One study reported that 11-month-old *Chchd2* deficient mice do not exhibit motor defects or OXPHOS impairment in the brain ^29^, while another research group has observed that older *Chchd2* knockout mice display mild motor and behavioral abnormalities and a subtle loss of midbrain dopamine (DA) neurons ^30^. Likewise, *CHCHD10*-deficient mice do not exhibit striking gross and molecular defects ^19^, suggesting that CHCHD2 and CHCHD10 may be functionally redundant and can partially compensate for each other. Despite the substantial interest in CHCHD2 and CHCHD10 in relation to human disease, numerous questions remain about the abundance, size, composition and function of the CHCHD2-CHCHD10 complex *in vivo*.

In this study, we generated a whole-body *Chchd2* knockout mouse to investigate the role of CHCHD2 in mitochondria. We thoroughly explored the consequences of CHCHD2 loss on motor performance and mitochondrial function in several tissues. Furthermore, we characterized the CHCHD2-CHCHD10 protein complex in mouse tissues and assessed its significance under conditions of mitochondrial dysfunction. Our findings provide new insights into the *in vivo* function of CHCHD2 in mammalian mitochondria and underscore its critical role in sustaining brain function and homeostasis.

## Results

### *Chchd2* ablation progressively impairs motor performance in male mice

To study the function of CHCHD2 in a mammalian living organism, we generated a mouse line carrying a conditional knockout allele by flanking exon 2 and 3 of *Chchd2* with loxP sites (Fig. S1A). The targeted allele was transmitted through the germline and heterozygous knockout mice (*Chchd2^+/-^*) were obtained by breeding *Chchd2^+/loxP^* mice with mice ubiquitously expressing cre-recombinase (b-actin cre). An intercross of heterozygous *Chchd2^+/−^* animals produced viable whole-body homozygous knockout (*Chchd2^−/−^*) pups, which were born at the expected Mendelian ratio (Fig. S1B). Western blot analyses confirmed that the CHCHD2 protein was absent in all investigated tissues from *Chchd2^−/−^* mice (Fig. S1C). Although these mice were fertile and appeared healthy in the first months of life, male knockouts were slightly smaller from 6 months of age (Fig. 1A) and presented a more prominent reduction in body weight at the age of 12 months (Fig. 1B). A similar reduction was observed in the weight of quadriceps (Fig. 1B), but no alterations were found in the size of the heart (Fig. 1B) or in organ-to-body weight ratios (Fig. S1D). In contrast, female *Chchd2^−/−^* mice did not exhibit decreased body weight at the age of 20 months (Fig. S1E).

**Fig. 1.**
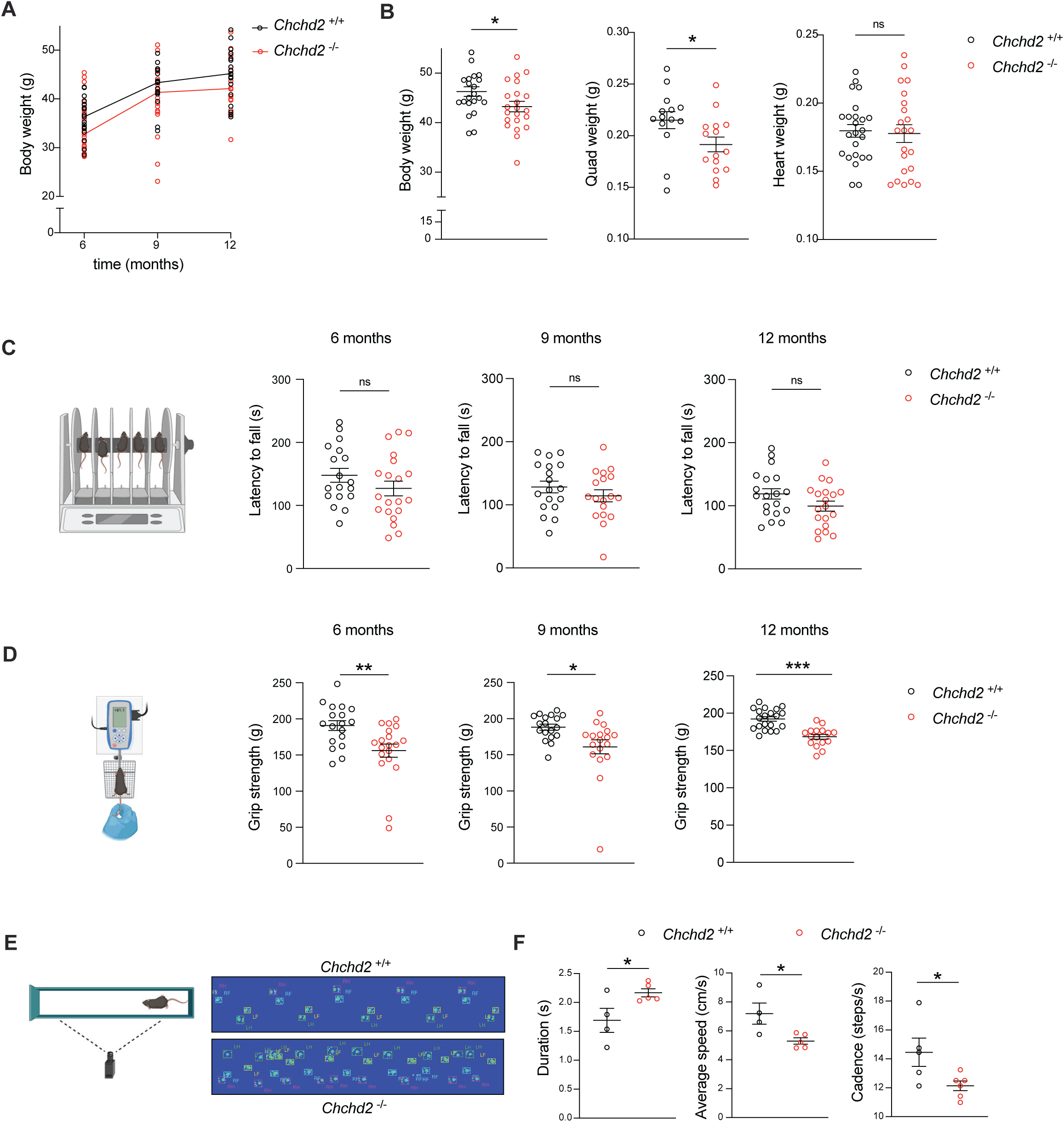
Male mice lacking CHCHD2 exhibit motor defects with age. **A.** Body weight (BW) of male *Chchd2* knockout (*Chchd2^−/−^)* and control (*Chchd2^+/+^)* mice at 6, 9 and 12 months; n>15 per genotype and time point. **B.** BW, quadriceps weight (QW) and heart weight (HW) measured in males at 12 months of age. Data are represented as means ± SEM; n>20 per genotype; *p < 0.05, ns not significant. **C.** Motor coordination measured as latency to fall (in s) from the rod in male mice from 6 to 12 months. **D.** Muscular strength (in g) measured by grip strength test in male mice from 6 to 12 months. Data are represented as means ± SEM *p < 0.05; n>18 per genotype. ns not significant. **E-F**. CatWalk gait analysis performed in males at 12 months of age. Representative digitized paw prints are shown in with RF, right front paw, RH, right hind paw, LF, left front paw, and LH, left hind paw. Duration (in s), average speed (cm/s) and cadence (steps/s) to cross the walkway in each step cycle). Data are presented as mean ± SEM. n≥ 4 *p < 0.05.

To define whether the loss of CHCHD2 alters animal motor performance, we tested spontaneous locomotion over time in open field arenas and found that vertical (rearing) and horizontal activities were unaffected in male mice lacking CHCHD2 (Fig. S2A). We next assessed motor coordination and muscular strength using rotarod (Fig. 1C) and grip (Fig. 1D) tests. These analyses revealed that male *Chchd2^−/−^* mice showed minor signs of impaired motor coordination and balance (Fig. 1C) and a significant reduction in muscular strength between the ages of 6 and 12 months (Fig. 1D). We performed gait analyses using the CatWalk apparatus and found that animals lacking CHCHD2 were slower and presented an unstable gait at 12 months (Fig. 1E, F). In contrast, female mice did not manifest abnormal motor function at the age of 20 months (Fig. S2B-D).

### Male mice lacking CHCHD2 present altered levels of monoamine neurotransmitters

We next asked whether the motor defects observed in male knockout mice were due to alterations in the levels of monoamine neurotransmitters, which modulate several physiological activities, including psychomotor function. We measured DA, norepinephrine (NE), epinephrine (EPI) and serotonin (5-HT) and their metabolites in different regions of the mouse brain using high-performance liquid chromatography (HPLC) (Fig.2A). At the age of 12 months, the striatal levels of DA and its precursor DOPA were significantly reduced in mice lacking CHCHD2 (Fig. 2B). A similar trend, although not significant, was observed in the DA metabolites 3,4-dihydroxyphenylacetic acid (DOPAC), homovanillic acid (HVA) and 3-methoxytyramine (3-MT) (Fig. 2B). NE and its metabolites 3-methoxy-4-hydroxyphenylglycol (MHPG) were reduced in the cerebellum and hippocampus (Fig. S3A, B), respectively, while EPI and the serotonin metabolite 5- hydroxyindole acetic acid (5-HIAA) were increased in the hippocampus (Fig. S3A). To follow up on the dysregulation of DA homeostasis found in the striatum, we assessed the striatal expression of the DA transporter (DAT) in the striatum. DAT levels were determined by autoradiography and no changes were observed upon CHCHD2 loss (Fig. 2C).

**Fig. 2.**
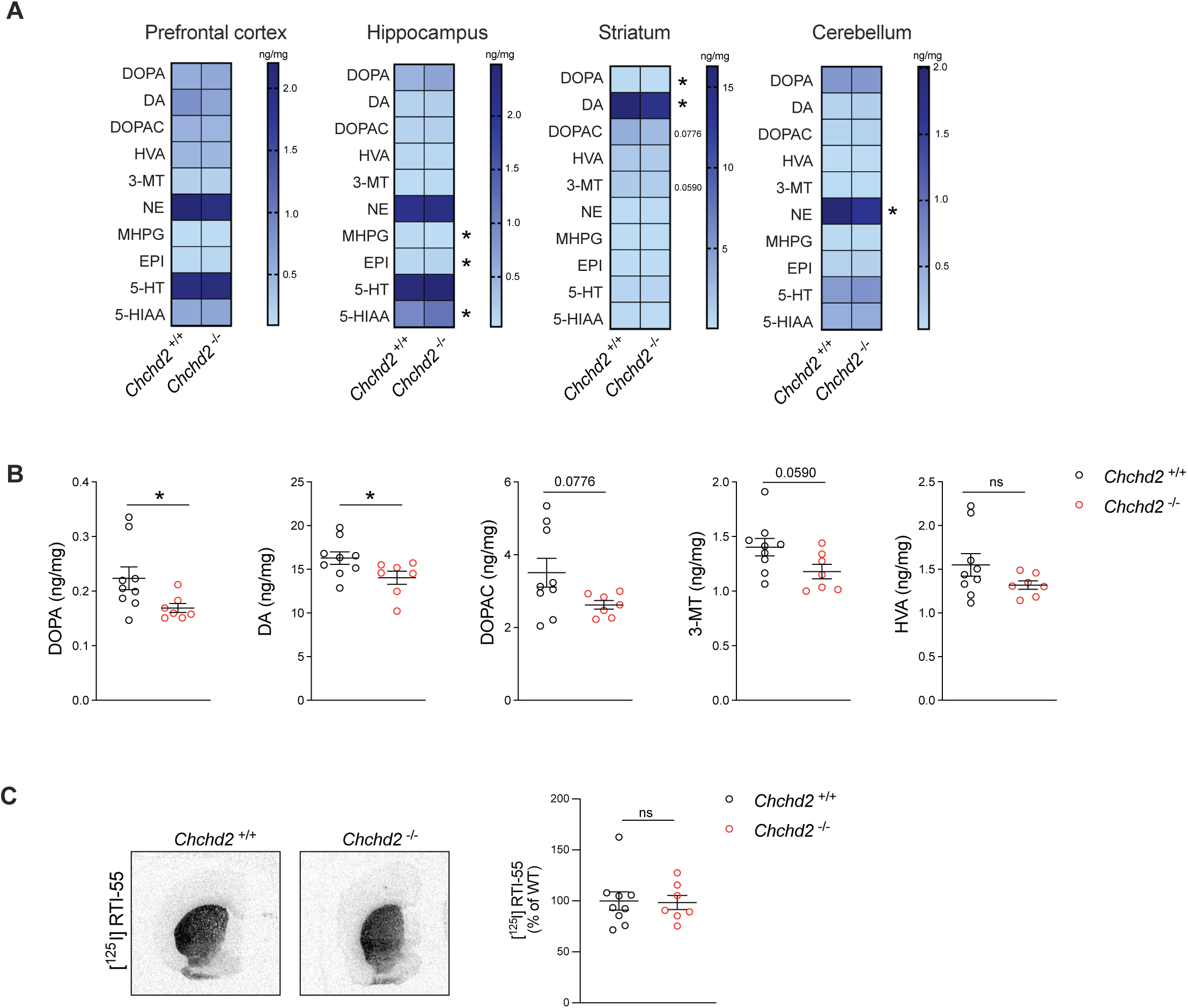
CHCHD2 ablation affects the levels of monoamine neurotransmitters in the brain of male mice. **A.** Heatmaps showing the levels (ng/mg) of DA, norepinephrine (NE) and serotonin (5-HT) and their metabolites in the prefrontal cortex, hippocampus, striatum, and cerebellum of *Chchd2^−/−^*and *Chchd2^+/+^* male mice at the age of 12 months. **B.** Levels (ng/mg) of DOPA, DA, 3,4-Dihydroxyphenylacetic acid (DOPAC), Homovanillic acid (HVA), 3-methoxytyramine (3-MT) in the striatum. Data are represented as means ± SEM; n≥ 7, *p < 0.05, ns not significant. **C.** Representative autoradiographs (left panel) and quantification (right panel) of [125I] RTI-55 binding with DAT on striatal sections of *Chchd2^−/−^* and *Chchd2^+/+^* male mice at the age of 12 months. Data are represented as means ± SEM; n≥ 5, ns not significant.

We proceeded to analyze the brain of 20-month-old females and found no significant changes in the prefrontal cortex, hippocampus, striatum, and cerebellum (Fig. S3C), in line with the lack of motor defects. However, the ratio of the HVA to DA in the prefrontal cortex was higher in *Chchd2^−/−^* females than in controls (Fig. S3D), which may reflect an increased DA turnover.

Taken together, our results demonstrate that male mice lacking CHCHD2 develop a mild phenotype in adult age, which manifests as a decline in body weight and muscle strength, gait abnormalities and DA depletion in the striatum. These findings align with a recent study showing that aged whole-body *Chchd2^−/−^* mice display PD-like phenotypic manifestations ^30^ and challenge the previously suggested hypothesis that CHCHD2 is dispensable for life ^14^.

### The absence of CHCHD2 does not reduce OXPHOS capacity but affects tissue morphology

We then examined the consequences of CHCHD2 ablation on mitochondrial function in several mouse tissues, including brain, heart and skeletal muscle. Western blot analysis at one year of age showed that *Chchd2^−/−^* mice had a mild decrease in steady-state levels of OXPHOS subunits in skeletal muscle and only a minor reduction in mitochondrially encoded cytochrome c oxidase I (MT-CO1) levels in the brain, whereas other tissues were unaffected (Fig. S4A). We assessed the activity of complex IV and complex V by using in-gel activity assays and found no alterations in any of the analyzed tissues (Fig. S4B). The activities of respiratory chain complexes, measured spectrophotometrically in mitochondria isolated from brain, heart and skeletal muscle, were also normal in *Chchd2^−/−^* mice at the age of one year (Fig. 3A). We additionally employed histochemical double staining to measure complex IV (cytochrome *c* oxidase, COX) and succinate dehydrogenase (SDH) activities *in situ*. These analyses revealed that in colon, a replicative tissue where mitochondrial damage can clonally expand with age ^31^, a few COX-deficient colonic crypts (blue cells) accumulated in response to CHCHD2 loss at the age of 12 months, while only one COX-deficient crypt was observed across different control animals (Fig. S4C).

**Fig. 3.**
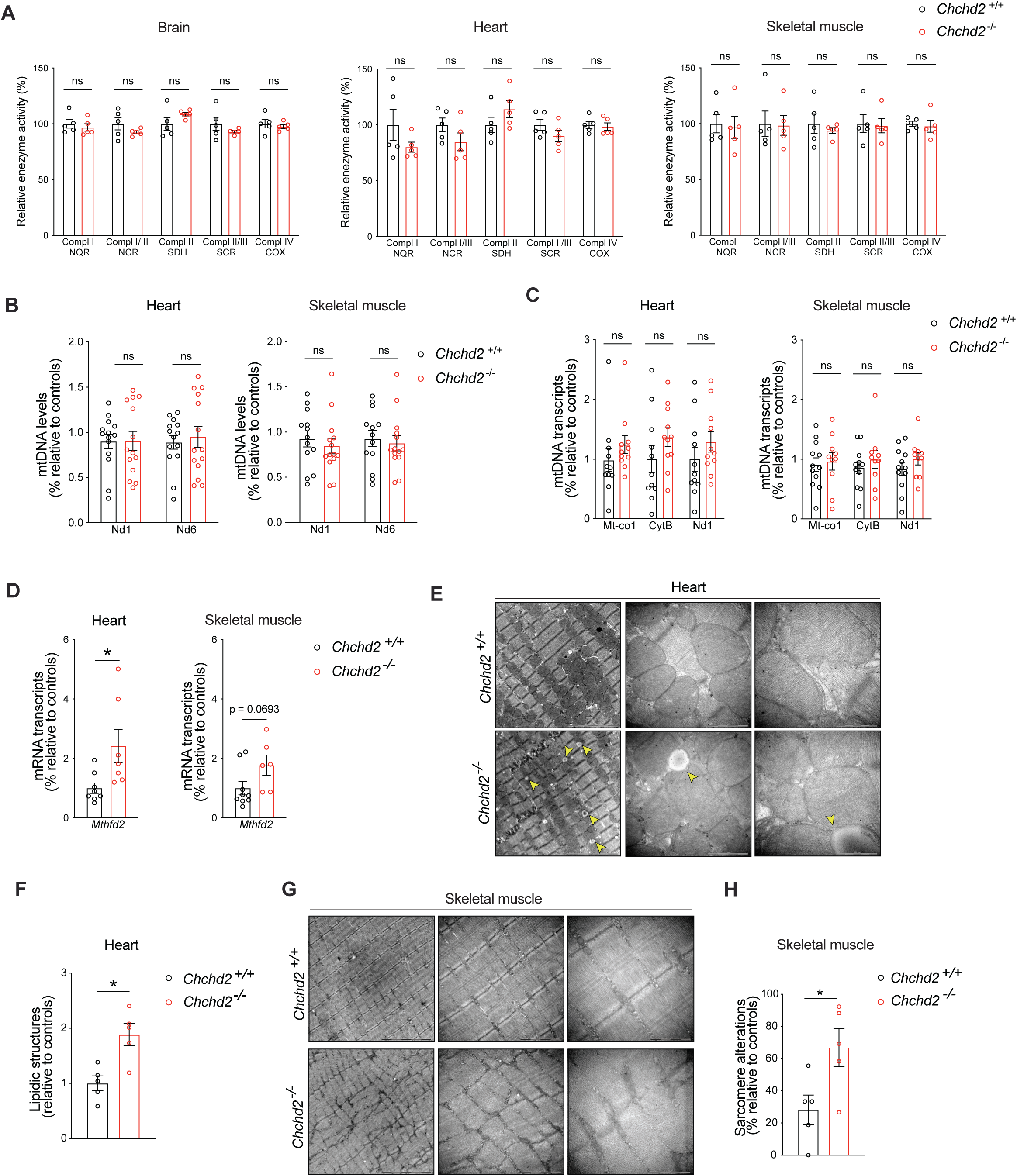
*Chchd2* knockout mice have normal OXPHOS activity but present mild alterations in tissue composition and morphology. **A.** Respiratory chain enzyme activity assays of CI, CI+CIII, CII, CII+CIII, and CIV in isolated mitochondria from brain, heart, and skeletal muscle (quadriceps) of *Chchd2^−/−^*and *Chchd2^+/+^* male mice at the age of 12 months. Data are represented as means ± SEM.; n=5; ns not significant. Quantification of **B.** mtDNA copy number (*Nd1* and *Nd6*/*18S rRNA*) and **C.** mtDNA transcripts (*Mt-Co1*, *CytB* and *Nd1*) performed by qPCR in brain, heart, and skeletal muscle. Data are represented as means ± SEM; *n* ≥ 11; ns not significant. **D.** *Mthfd2* gene expression measured by qPCR in heart and skeletal muscle of male mice at 12 months of age. Data are represented as means ± SEM; *n* ≥ 11; *p< 0.05; **E-F.** Representative electron microscopy images of tissue and mitochondrial morphology (Scale bar: 5, 1 μm and 500 nm) and sample-wise quantification of vacuoles in the heart of male mice at 12 months of age. **G-H.** Representative electron microscopy images of tissue and mitochondrial morphology (Scale bar: 2 and 1 μm) and **s**ample-wise quantification of sarcomere alterations in the skeletal muscle of male mice at the age of 12 months. Data are represented as means ± SEM; *n*=5; * p< 0.05.

Given that mutations in *CHCHD10* were reported to cause accumulation of mtDNA deletions and reduction in mtDNA levels in both human and mouse tissues^14,20,32^, we measured mtDNA copy number in heart and skeletal muscle by qPCR. No changes were found when comparing control and knockout mice at the age of one year (Fig. 3B). Moreover, the levels of mtDNA transcripts in these tissues were unaltered (Fig. 3C), suggesting that CHCHD2 is not involved in the maintenance or integrity of mtDNA gene expression. However, the heart and skeletal muscle of *Chchd2* knockout animals were characterized by an upregulation (~1.8- to 2.5-fold) of the methylenetetrahydrofolate dehydrogenase 2 (*Mthfd2*) gene, encoding a key enzyme in mitochondrial one-carbon metabolism pathway that serves as an early marker for mitochondrial dysfunction ^33^ (Fig. 3D). Interestingly, *Mthfd2* expression did not correlate with the activation of mitochondrial integrated stress response (ISR^mt^), as shown by the normal transcript levels of *Atf4*, *Atf5*, *Chop* and *Fgf21* (Fig. S4D).

We thereafter investigated whether the loss of CHCHD2 causes any disruptions in tissue morphology or mitochondrial cristae organization. Electron microscopy analysis of one-year-old mice identified a higher content of circular vacuoles resembling lipid droplets in the heart of *Chchd2* knockout mice (Fig. 3E, F). Similar vacuoles were previously described in cardiac and skeletal muscle of mice expressing pathological mutants of CHCHD10^34^ and in the skeletal muscle of patients carrying CHCHD10 mutations causing myopathy^32^. Electron micrographs of *quadriceps femoris* demonstrated the presence of alterations in the sarcomere organization with irregular arrangement of myofibrils and the z-discs in longitudinal sections of mice lacking CHCHD2 (Fig. 3G, H). However, mitochondrial ultrastructure and cristae organization were preserved in heart and skeletal muscle (Fig. 3G, H). Based on these findings, we conclude that CHCHD2 is not required to sustain OXPHOS capacity in mouse tissues under physiological conditions, but it is involved in preserving tissue morphology and lipid content.

### Loss of CHCHD2 alters lipid homeostasis in the cortex and dorsal striatum

The identification of lipid vacuoles in the heart of mice lacking CHCHD2 prompted us to assess the lipid levels in the brain of our mouse model. Given that the brain comprises approximately 50% dry mass of lipids^35^, which are vital for both its structure and function, we performed an untargeted spatial lipidome analysis using high-throughput dual polarity matrix-assisted laser desorption/ionization (MALDI)-FTICR-MSI method ^36,37^. This analysis revealed significant alterations in the cortex and dorsal and ventral striatum of *Chchd2*-deficient mice at 12 months of age (Fig. 4A, B, Fig. S5A and Listed in Supplementary table 6). Non-hydroxylated sphingolipids (d=nonhydroxylated), such as sulfohexosylceramides (SHexCers), hexosylceramides (HexCers) and sphingomyelins (SMs) were increased in dorsal striatum and cortex, while the hydroxylated SHexCers (t=hydroxylated) were decreased in the cortex (Fig. S5A). Notably, similar trends in the levels of hydroxylated and non-hydroxylated sulfatides were recently found in the brain tissue sections of non-human primates treated with 1-methyl-4-phenyl-1,2,3,6-tetrahydropyridine (MPTP), which is widely used to generate PD models ^38^. We found additional changes in different species of phospholipids, such as phosphatidylinositol (PI), phosphatidylserines (PS) and phosphatidylcholine (PC), which are essential components of mammalian membranes including mitochondrial membranes. The polyunsaturated fatty acid-containing PI (40:6) and PI (38:6) were higher in the cortex of *Chchd2* knockout mice. In contrast, the phosphatidylserines PS (40:4) and PS (40:6) were decreased in the cortex and dorsal striatum, respectively. The levels of the phosphatidylcholine species PC (40:4) and PC (30:0) were also affected in these two brain regions (Fig. S5A). Notably, cardiolipins, unique phospholipid components of mitochondrial membranes were unchanged upon CHCHD2 loss (Data not shown). Overall, the loss of CHCHD2 causes alterations in lipid homeostasis in different brain regions, including the dorsal striatum, which is particularly affected in patients with PD and where a significant reduction in DA levels was found.

**Fig. 4.**
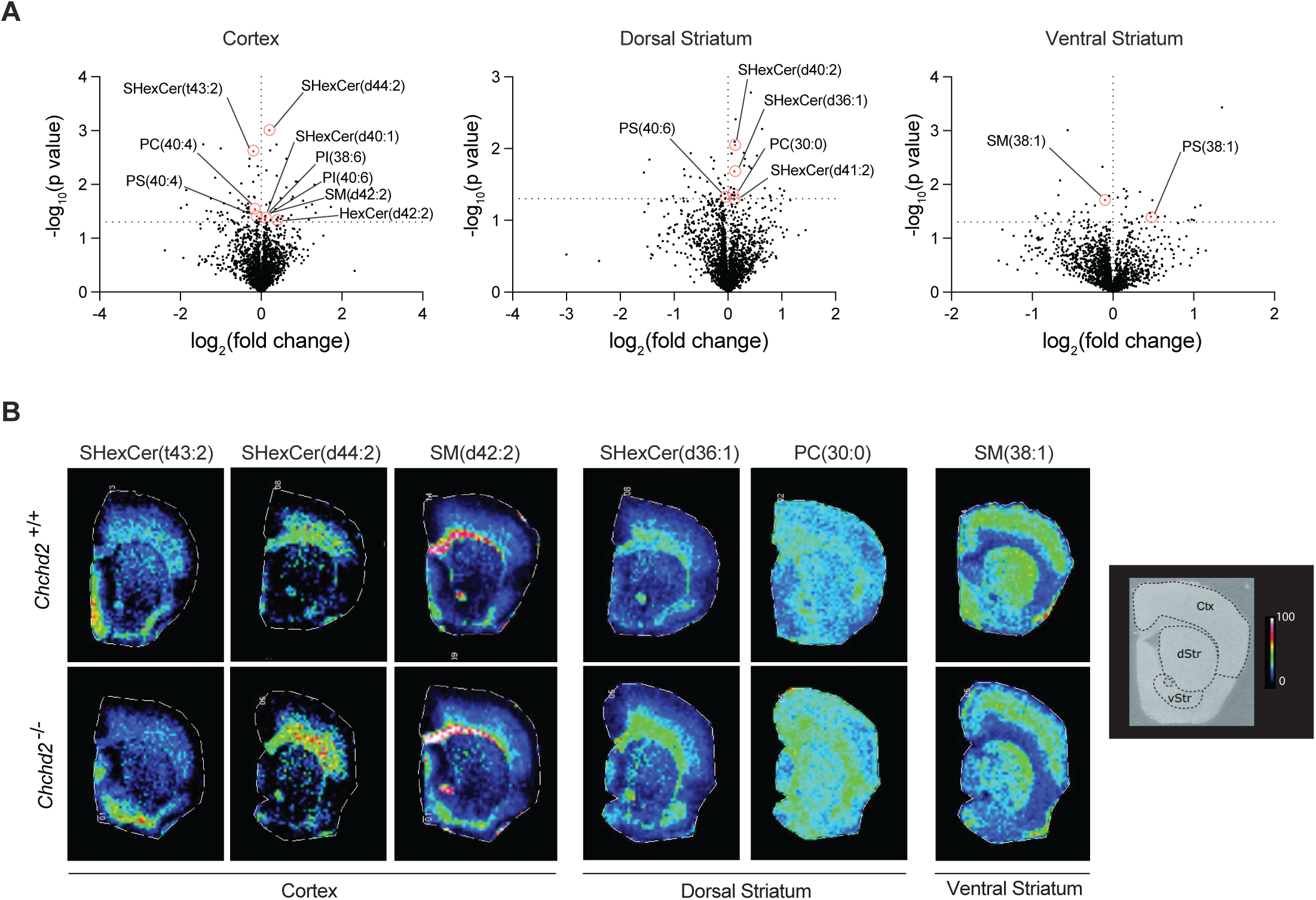
Male mice lacking CHCHD2 exhibit regional alterations in lipid content in the brain. **A.** Volcano plots showing log_2_ (fold changes) and the −log_10_ (p values) of all detected *m/z* values in the cortex, dorsal striatum and ventral striatum in *Chchd2^−/−^*and *Chchd2^+/+^* male mice at the age of 12 months. Identified regulated lipid are highlighted with the monoisotopic peaks of their corresponding ions with red dash lines. **B.** Representative ion images of SHexCers (sulfohexosylceramides), HexCers (hexosylceramides), SMs (sphingomyelins), and PCs (Phosphatidylcholines) peaks in the cortex, dorsal and ventral striatum of *Chchd2^−/−^*and *Chchd2^+/+^* male mice at the age of 12 months. The lateral resolution of the MALDI-FITCR-MS ion images is 100 µm, and all of the ion images were RMS-normalized.

### *In vivo* CHCHD2 and CHCHD10 are fully assembled into a large protein complex

Previous studies performed in human cell cultures have shown that CHCHD2 and CHCHD10 are highly abundant as monomers or dimers, but they can also form high molecular weight complexes of ~140-220 kDa^18,19^. To investigate the abundance and the size of this protein complex *in vivo*, we performed two-dimensional gel electrophoresis (2D-PAGE) analyses in mitochondria isolated from brain, heart and skeletal muscle of *Chchd2^+/+^* and *Chchd2^−/−^* mice using either mild (digitonin) (Fig. S6A) or stringent (DDM) detergent solubilization conditions (Fig. 5A). These analyses revealed that in all analyzed tissues from control mice CHCHD2 and CHCHD10 co-migrated only as a high molecular complex of ~140 kDa and no monomers or dimers were found (Fig. 5A). Remarkably, in the absence of CHCHD2, CHCHD10 could still oligomerize, although it formed a less abundant, possibly unstable complex (Fig. 5A). Moreover, neither CHCHD2 nor CHCHD10 co-migrated with CHCHD3, indicating that these two proteins are not MICOS subunits. To confirm that the observed differences in the oligomerization capacity of CHCHD2 and CHCHD10 were not related to different experimental conditions, we performed 2D-PAGE using mitochondria isolated from human HEK293T cells and primary mouse embryonic fibroblasts (MEFs) (Fig. S6B, C). In these tissue culture cells, the large majority of CHCHD2 and CHCHD10 were found at low molecular weight, presumably as monomers or dimers, although a portion of CHCHD2 migrated at higher molecular weight. To characterize the CHCHD2-CHCHD10 complex at higher resolution, we employed complexome profiling proteomics of skeletal muscle mitochondria resolved by BN-PAGE (Fig. S6D). Analysis of the electrophoretic migration profiles showed a striking comigration between CHCHD2 and CHCHD10 at molecular weights ranging from 100 to 140 kDa (fractions 28-45) (Fig. 5B). In agreement with our 2D gels results, CHCHD2 and CHCHD10 were not detected in low molecular weight fractions (fractions 50 to 64), excluding their existence as monomers or dimers *in vivo* (Fig. 5B). In addition, we found that the migration profile of CHCHD10 was not significantly affected in *Chchd2^−/−^* mice (Fig. S6E), validating that CHCHD10 is still able to form high molecular weight oligomers upon CHCHD2 loss. Notably, the absence of CHCHD2 did not affect the stability or the assembly of MICOS (Fig. S6F).

**Figure 5.**
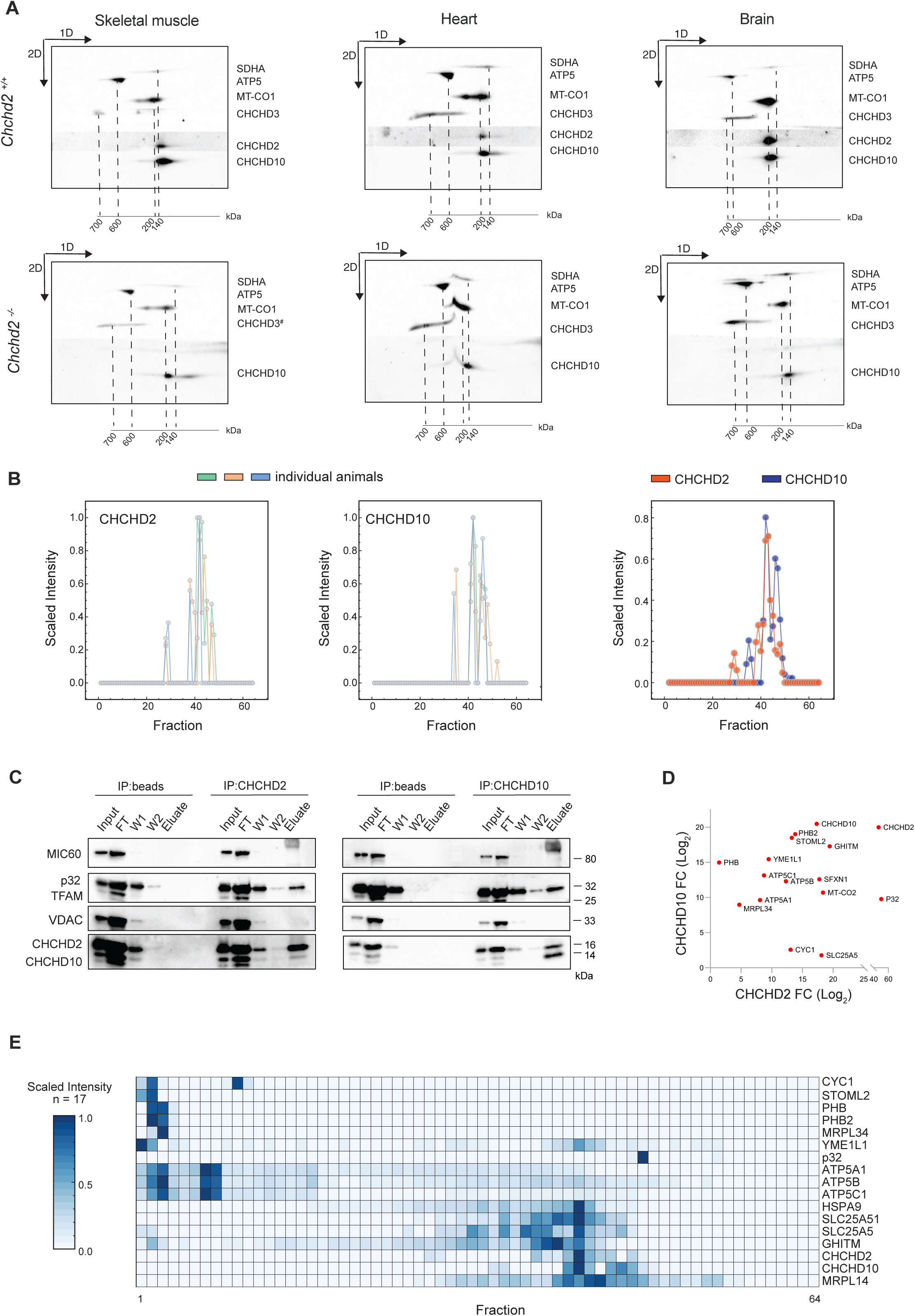
*In vivo* CHCHD2 and CHCHD10 are monomers but are fully assembled into a large complex. **A.** 2D-PAGE analyses of mitochondria isolated from skeletal muscle, heart and brain of Chchd2^−/−^ and Chchd2^+/+^ male mice at the age of 12 months. Mitochondria were solubilized using 1% (w/v) DDM. The position of SDHA, ATP5, MT-CO1, and CHCHD3 corresponds to the size of their different protein complexes (in kDa). The molecular weights of monomeric CHCHD2 and CHCHD10 are 15.5 and 14.1 kDa, respectively. **B.** Migration profiles of CHCHD2 and CHCHD10 obtained from complexome profiling analysis of mitochondria isolated from skeletal muscle of *Chchd2^+/+^* mice at the age of 12 months. n=3. **C.** Western Blot analysis of CHCHD2 and CHCHD10 pull-down experiments performed using mitochondria isolated from HEK293T cells. ab=antibody; FT=flow through; W=wash. **D.** Interactome plot of proteins identified by CHCHD2 and CHCHD10-IP. Only significant hits (p<0.05) are visualized, and axes represent respective Log_2_FC of each protein between IP pull-down samples and controls. **E.** Heatmap obtained from complexome analysis of skeletal muscle mitochondria (n = 3) and depicting the relative abundances of proteins found significantly enriched in the eluate of CHCHD2 and CHCHD10 pulldowns at FDR level of 0.05.

Next, we investigated the composition of the complex by co-immunoprecipitation (co-IP) experiments of the endogenous proteins using mitochondria isolated from wild-type HEK293T cells. Western blot analysis confirmed the strong physical interaction between CHCHD2 and CHCHD10, which were successfully co-immunoprecipitated using specific antibodies (Fig. 5C). The previously reported interacting partner p32 (C1QBP), was also found in both CHCHD2 and CHCHD10 eluates (Fig. 5C). Label-free quantitative mass spectrometry was applied to the pulldown eluates to determine additional interacting partners (Fig. S6G, H and Listed in Supplementary table 3, 4). This analysis revealed that CHCHD2 and CHCHD10 interactomes largely overlapped (Fig. 5D and Supplementary table 5). We observed that stomatin-like protein 2 (SLP2) and the iAAA protease YME1L were highly enriched in both CHCHD2 and CHCHD10 pulldowns (Fig. 5D), while the rhomboid protease PARL was only found as a CHCHD10 interacting partner (Fig. S6H). In line with our results, it was previously reported that SLP2-PARL-YME1L form a large protease complex, known as SPY, which interacts with CHCHD2 ^39^. Furthermore, both subunits of prohibitin (PHB) complex, PHB1 and PHB2, were co-immunoprecipitated with CHCHD2 and CHCHD10. Interestingly, a recent study in mutant mice carrying a *Chchd10* pathological variant has shown that CHCHD10 together with the protease SLP2 controls the stability of the PHB complex^27^.

To discriminate whether the identified proteins were merely interacting partners or additional subunits of the CHCHD2-CHCHD10 protein complex, we surveyed our complexome profiling datasets and generated a heatmap of their relative abundance in the different fractions (Fig. 5E). We found that none of the molecular interactors, including p32, presented a pattern fully matching CHCHD2 and CHCHD10 migration profiles, suggesting that these proteins are not components of the complex.

Altogether, our results demonstrate that CHCHD2 and CHCHD10 do not exist as free monomers *in vivo* but are fully assembled into a complex of about 100-140 kDa. In contrast to a recent report ^40^, we show that despite its strong physical interaction, p32 is not a scaffold subunit of the CHCHD2-CHCHD10 complex. Finally, we demonstrate that in mouse tissues, the loss of CHCHD2 does not prevent the formation of CHCHD10 oligomers, challenging the hypothesis that CHCHD2 is required for CHCHD10 oligomerization ^41^.

### The levels and the size of the CHCHD2-CHCHD10 complex are regulated by the severity of mitochondrial dysfunction

Recently, it was reported that CHCHD2 and CHCHD10 accumulate in mitochondria of human cell lines treated with the protonophore m-chlorophenylhydrazone (CCCP)^41^. Additionally, the *Drosophila* ortholog of CHCHD2 was found upregulated under mitochondrial stress conditions^28^. These data suggest that CHCHD2 and CHCHD10 expression or stability is influenced by mitochondrial damage and cellular metabolism. To test this hypothesis, we analyzed CHCHD2 and CHCHD10 protein levels in a variety of *in vivo* and *in vitro* models affected by severe mitochondrial dysfunction (Fig. 6A). We first interrogated publicly available proteomic datasets of heart conditional knockout mouse strains characterized by defects in mtDNA replication, maintenance, transcription and translation caused by ablation of the mitochondrial DNA helicase (*Twinkle*), the mitochondrial transcription factor A (*Tfam*), the mitochondrial transcription elongation factor (*Tefm*) and the mitochondrial transcription termination factor 4 (*Mterf4),* respectively ^33,42^. At a late-disease stage, when the levels of MTHFD2 had increased by 70- to 215-fold, the CHCHD2 and CHCHD10 were significantly upregulated by approximately 1.5- to 2.5-fold and 4- to 10-fold in the hearts of these different knockouts (Fig. 6B). A significant increase in CHCHD2 and CHCHD10 was also observed when analyzing the proteomes of human cell lines lacking accessory subunits of complex I ^43^ (Fig. 6B). These findings led us to hypothesize that the upregulation of CHCHD2 and CHCHD10 might be a common response to mitochondrial damage across different species, organs and tissues. Hence, we experimentally examined different tissues from four mouse models with defective mitochondrial function at their respective experimental endpoints. We analyzed the heart of 24-week-old *Ribonuclease H*1 (*RNaseH1*) conditional knockouts ^44^, the skeletal muscle of 21-week-old *Tfam* conditional knockout mice ^45^, and the liver of wild-type mice treated with the inhibitor of mitochondrial transcription (IMT) for four weeks ^46^. As a positive control, we included the heart of the previously analyzed 8-week-old *Tefm* knockout model ^42^. We initially determined the magnitude of mitochondrial impairment by measuring the levels of OXPHOS subunits (Fig. S7A) and complex I and complex IV activities (Fig. 6C and Fig. S7B), which demonstrated a dramatic reduction in OXPHOS capacity in each tissue and mouse model. We additionally measured the expression of *Mthfd2*, which is transcriptionally regulated and correlates with severity and progression of mitochondrial defects ^47^. We found that *Mthfd2* transcript levels were strongly upregulated (~80 to 150-fold) in heart and skeletal muscle of mice lacking RNAseH1, TEFM and TFAM (Fig. S7C), in line with severe mitochondrial damage leading to lethality^33^. In the liver of IMT-treated mice, the increase in *Mthfd2* transcript levels was much milder (~5-fold) (Fig. S7C), consistent with no effects on animal survival and organ function ^46^. Next, we assessed the steady-state levels of CHCHD2 and CHCHD10 by western blot analysis using mitochondria isolated from heart, skeletal muscle and liver (Fig. 6D and Fig. S7D). We found that the CHCHD10 protein levels were markedly increased in heart and skeletal muscle of mice lacking RNaseH1, TEFM and TFAM, whereas it was only moderately increased in the liver of IMT-treated mice (Fig. 6D and Fig. S7D). Although less pronounced, CHCHD2 was also upregulated in all the models and tissues (Fig. 6D and Fig. S7D), in line with the results obtained from the proteomics datasets. Next, we studied whether changes in gene expression were responsible for CHCHD2 and CHCHD10 increase. We thus measured mRNA levels by RT-qPCR (Fig. 6E and Fig. S7E) and found that *Chchd2* transcripts were unchanged across all tissues and animal models. In contrast, *Chchd10* mRNA levels were mildly upregulated in the heart of *Tefm* knockout mice (~2.5-fold) and the liver of IMT-treated mice (~1.5-fold) (Fig. 6E and Fig. S7E). A similar trend, although not significant, was observed in the heart of *RNaseH1* knockout mice and the skeletal muscle of *Tfam* knockout mice (Fig. 6E). However, the upregulation of *Chchd10* transcription was not as pronounced as the increase in protein levels.

**Fig. 6.**
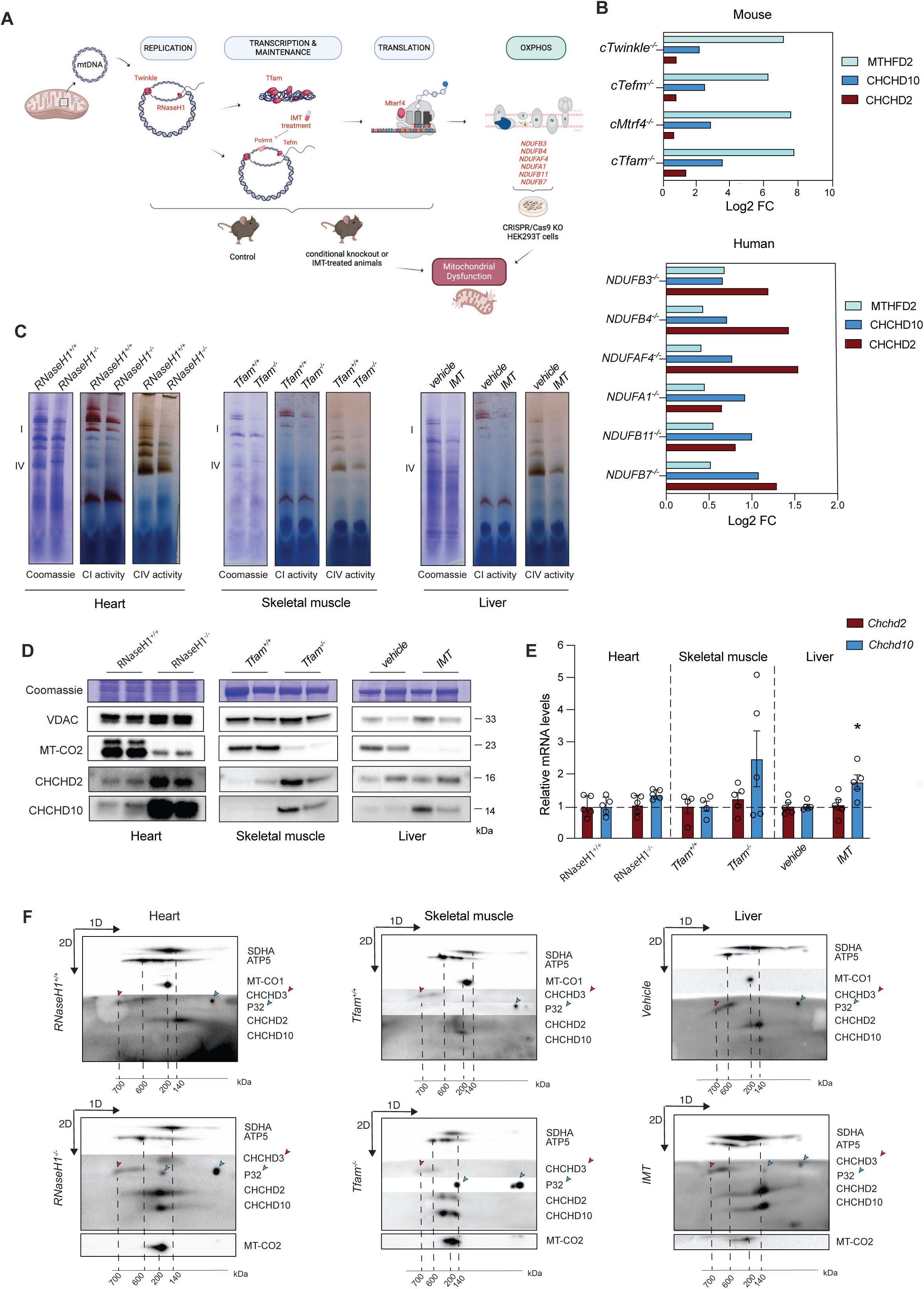
The levels and the size of the CHCHD2-CHCHD10 protein complex are modulated in response to mitochondrial damage. **A**. Schematic representation of models with mitochondrial dysfunction. The image was created with BioRender.com. **B-C**. Levels of MTHFD2, CHCHD2, and CHCHD10 (Log_2_FC) obtained using publicly available proteomic datasets from tissue-specific knockout mouse strains with impaired mtDNA gene expression (*Twinkle, Tefm, Mterf4* and *Tfam* knockout mice) and in HEK293T cell lines lacking complex I accessory subunits. **D.** Complex IV and V in-gel activities performed on mitochondria isolated from heart tissue of *RNaseH1* mice at 24 weeks of age, skeletal muscle (gastrocnemius) of *Tfam* mice at 5 months of age and liver of IMT treated mice for four weeks and respective controls. BN-PAGE were stained with Coomassie or incubated with substrates for detecting the in-gel activity of the indicated OXPHOS complexes. **E.** Western Blot analysis of CHCHCD2 and CHCHD10 steady-state levels in mitochondria isolated from tissues of three different mouse strains affected by mitochondrial dysfunction in heart, skeletal muscle and liver, respectively. VDAC and Coomassie staining were used as loading controls. **E.** *Chchd2* and *Chchd10* mRNA levels measured by qPCR from RNA isolated from mouse tissues with mitochondrial dysfunction. Data are represented as mean ± SEM. n= 5 mice for each genotype, *p < 0.05. **F.** 2D-PAGE of mitochondria isolated from skeletal muscle of from heart tissue of *RNaseH1* mice at 24 weeks of age, skeletal muscle (gastrocnemius) of *Tfam* mice at 5 months of age and liver of IMT treated mice for four weeks and respective controls Mitochondria were solubilized using 1% (w/v) DDM. The position of SDHA, ATP5, MT-CO1, MT-CO2 (used in models of mitochondrial dysfunction) and CHCHD3 (indicated by a red arrow) corresponds to the size of their different protein complexes (in kDa). P32 is indicated by blue arrows.

We next determined whether the increase in CHCHD2 and CHCHD10 protein levels in response to mitochondrial impairment is reflected in the complex formation, we employed 2D-PAGE on mitochondria isolated from heart, skeletal muscle and liver from each mouse strain (Fig. 6F and Fig. S7F). Consistent with our previous results, CHCHD2 and CHCHD10 run as a complex of ~140 kDa in healthy mitochondria. In defective mitochondria, the abundance of the CHCHD2-CHCHD10 complex was markedly increased without the accumulation of any smaller forms. Surprisingly, the CHCHD2-CHCHD10 complex presented a notable shift towards higher molecular weight upon mitochondrial defects in heart, skeletal muscle and liver (Fig. 6F and Fig. S7F). Furthermore, mitochondrial dysfunction also affected the migration pattern of a fraction of p32, which co-migrated with CHCHD2 and CHCHD10, suggesting that in response to mitochondrial damage p32 may either oligomerize or structurally contribute to the formation of the CHCHD2-CHCHD10 complex.

Taken together, our results indicate that the increase in CHCHD2 and CHCHD10 levels is a common response across different tissues affected by mitochondrial dysfunction. This upregulation is proportional to the severity of mitochondrial damage and corresponds to an increase in both the abundance and size of the CHCHD2-CHCHD10 complex.

### CHCHD2 upregulation protects against mitochondrial damage

To investigate whether the upregulation of the CHCHD2-CHCHD10 complex leads to a selective advantage in damaged mitochondria, we assessed the growth and survival rate of primary mouse embryonic fibroblasts (MEFs) under different stress conditions. We hence generated and analyzed four *Chchd2* knockout and four control MEF lines (Fig. 7A). The loss of CHCHD2 caused ~50% decrease in the mtDNA-encoded OXPHOS subunit MT-COI in primary MEFs (Fig. 7B) and a significant reduction (~50%) in cell proliferation in galactose medium, where cells were forced to rely on OXPHOS for ATP production (Fig. 7C). We next measured how treatment with the complex I inhibitor rotenone or the antibiotic actinonin, which compromises mitochondrial translation, impacts CHCHD2 and CHCHD10 levels. Western blot analyses showed that CHCHD2 and CHCHD10 were upregulated 24 hours after rotenone treatment (Fig. S8A), whereas the exposure to actinonin caused a clear accumulation of CHCHD2 and CHCHD10 only after 72 hours (Fig. S8B). Based on these observations, we analyzed *Chchd2* knockout and control MEFs exposed to rotenone for 24 to 72 hours and actinonin for 72 to 118 hours. Importantly, we found that the ablation of CHCHD2 did not prevent CHCHD10 accumulation induced by these mitochondrial insults, but it rather enhanced it (Fig. 7D, E), suggesting that the upregulation of these proteins is beneficial to counteract the mitochondrial impairment. To test this hypothesis, we measured cell viability after rotenone and actinonin treatments and found that both agents caused a substantial decrease in cell viability in control MEFs (Fig.7F, G). Notably, cells lacking CHCHD2 were significantly more sensitive to rotenone and actinonin (Fig.7F, G).

**Fig. 7.**
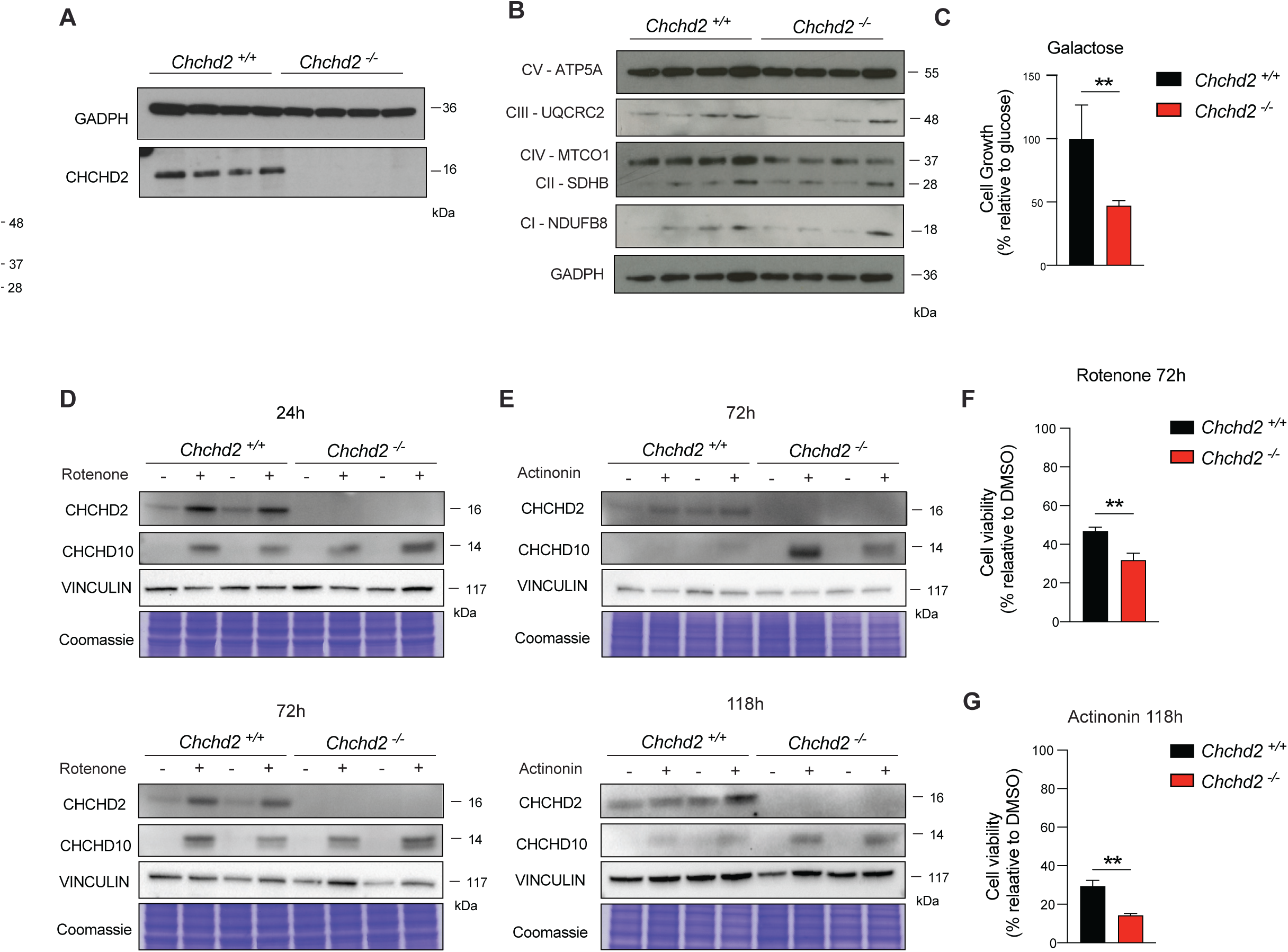
CHCHD2 is required to sustain cell viability under mitochondrial stress conditions. **A-B.** Western blot analyses of steady-state levels of CHCHD2 and OXPHOS subunits in total extracts from primary *Chchd2^−/−^* and *Chchd2^+/+^* MEFs (n=4 cell lines per genotype). GADPH was used as a loading control. **C.** Analysis of cell growth in medium containing 0.9 g/l galactose. Data are expressed as % relative to the cell growth in medium containing glucose and represented as mean ± SEM (n=4 cell lines per genotype); **p < 0.01. **C-D.** Representative Western Blot analysis of CHCHCD2 and CHCHD10 steady-state levels in total extract from primary *Chchd2^−/−^* and *Chchd2^+/+^* MEFs treated with 250 nM rotenone for 24 and 72 hours and 150 nM actinonin for 72 and 118 hours. VINCULIN and Coomassie were used as loading control. **E-F.** Cell viability performed in primary *Chchd2^−/−^* and *Chchd2^+/+^* MEFs treated with 250 nM rotenone for 72 hours and 150 nM actinonin for 118 hours. Data are represented as means ± SEM; *n*=4 per genotype; **p < 0.01.

Overall, our *in vivo* and *in vitro* results indicate that CHCHD2 is involved in cellular responses to mitochondrial damage. These data argue that CHCHD2 upregulation provides protection against mitochondrial dysfunction and CHCHD10 can only partially compensate for CHCHD2 loss.

## Discussion

Several mutations in *CHCHD2* and *CHCHD10* genes have been found in patients with neurodegenerative diseases. However, it remains unclear whether the pathophysiology is driven by gain-of-function or loss-of-function mechanisms. Research efforts have largely focused on understanding the impact of mutations causing the accumulation of toxic species, whereas the physiological function of CHCHD2 and CHCHD10 in the IMS remains poorly understood. This study explores the role of CHCHD2 using mammalian systems and examines the consequences of its loss on mouse gross and molecular phenotypes. We here show that adult male mice lacking CHCHD2 manifest a modest decrease in muscle strength and gait abnormalities, likely caused by the decreased DA levels in the striatum. These findings suggest that the loss of *CHCHD2* does not compromise the early development of the nigrostriatal system. In line with this hypothesis, single-cell RNAseq data have shown that mouse ventral midbrain neurons express *CHCHD2* and *CHCHD10* at low levels during embryonic and perinatal stages while both genes are strongly upregulated in adulthood ^48^. This temporal expression pattern may explain why *CHCHD2* loss does not affect juvenile mice but leads to defects in adult age, offering a plausible explanation for the onset of neurodegenerative disorders in humans.

Although CHCHD2 was first found as a top candidate in a computational screen for novel regulators of OXPHOS function ^49^, it remains unclear whether the lack of this protein affects the respiratory capacity. Previous studies have reported that CHCHD2 can regulate complex IV activity acting as a transcription factor of the COX4I2 subunit in cells under stress conditions ^50^, however, the nuclear localization of CHCHD2 is largely debated. In the yeast *Saccharomyces cerevisiae,* the disruption of the CHCHD2/CHCHD10 ortholog, Mix17, leads to a mild reduction in oxygen consumption and lower activity in complex III, while complex IV activity is maintained^51^. In *Drosophila,* CHCHD2 disruption mildly reduces ATP levels in the thorax muscle ^28^, while knocking out *Chchd2* in zebrafish compromises the assembly of Complex I ^52^. Additional studies reported that CHCHD2 ablation causes a profound decrease in basal and maximal oxygen consumption in glioblastoma cells ^53^. Conversely, HEK293T cells lacking CHCHD2 maintain normal respiratory complex activities ^41^. Overall, there is still no clear consensus on whether CHCHD2 loss affects OXPHOS capacity. Here, we assessed the activities of OXPHOS complexes across various tissues in CHCHD2-deficient mice at one year of age. Our data demonstrate that loss of CHCHD2 does not impair mitochondrial respiration in differentiated tissues, although marginally increases the number of COX-deficient cells found in the highly regenerative colonic epithelium. These findings indicate that CHCHD2 is dispensable for tissue bioenergetics in physiological conditions.

Mutations in *CHCHD2* and *CHCHD10* have been predominantly associated with cristae abnormalities ^26,54^. Although these alterations were initially attributed to the destabilization of the MICOS complex ^26^, more recent investigations suggest that the abnormal mitochondrial ultrastructure linked to CHCHD2 and CHCHD10 mutations is caused by an excessive cleavage of the mitochondrial optic atrophy 1 (OPA1), a key component of the mitochondrial fusion machinery^34,55^. However, the hearts of one-year-old CHCHD2-knockout mice do not exhibit any disruptions in cristae organization but present a clear accumulation of lipid vacuoles. Additionally, the brains of these animals showed altered lipid content, with significant changes observed in myelin-associated sphingolipid species within the cortex and dorsal striatum, whereas few phospholipid species were dysregulated in these regions. In line with our observation, a recent study reported the presence of phospholipid alterations in the heart of CHCHD10 mutant mice ^57^. Given that lipids serve as signaling molecules and structural components of cellular membranes, it is plausible that these disruptions may contribute to the mitochondrial damage seen with CHCHD2 mutations. It is also noteworthy that another member of the CHCH-domain protein family, Mdm35 (TRIAP in humans), is involved in the regulation of lipid content by mediating the transfer of phospholipids between the outer and inner mitochondrial membranes ^58,59^. Although the underlying molecular mechanisms remain to be elucidated, these data suggest that CHCHD2 is involved in maintaining lipid homeostasis.

In proliferative cells, CHCHD2 and CHCHD10 are mainly present as monomers or dimers^18,19^. A similar distribution has been observed for CHCHD10 in zebrafish larvae ^52^, raising questions about the functional relevance of the CHCHD2-CHCHD10 protein complex. We here provide compelling evidence that in differentiated mouse tissues, such as brain, heart and skeletal muscle, CHCHD2 and CHCHD10 exist exclusively as a complex of ~100-140 kDa, suggesting that these proteins do not function as free monomers. In contrast to earlier reports ^41,60^, we show that CHCHD10 can oligomerize in the absence of CHCHD2, which may explain how these functionally redundant proteins can compensate for one another. The analysis of CHCHD2 and CHCHD10 interactomes did not uncover any additional components of the complex but confirmed significant interactions with the mitochondrial proteolytic system within the IMS and matrix, including the SPY complex, PHB complex, and GHITM (also known as TMBIM5 or MICS1). These interactions may either point to a role in maintaining mitochondrial proteostasis or could account for the rapid turnover of CHCHD2 and CHCHD10. Notably, their half-lives in cell lines range from 2 to 5 hours ^19,61^, while the median protein half-life in mitochondria is about 87 hours ^61^. This implies that CHCHD2 and CHCHD10 are constantly degraded and thus their levels are finely tuned under physiological conditions. Meanwhile, in response to mitochondrial stress CHCHD2 and CHCHD10 accumulate, likely due to increased protein stability ^28,41^. Our data reveal that this process is highly conserved *in vivo* and represents a common response across various tissues and organs affected by mitochondrial dysfunction. Strikingly, the increase in CHCHD2 and CHCHD10 correlates with the severity of mitochondrial defects, leading to the formation of larger and more abundant protein complexes. In MEFs, the absence of CHCHD2 does not prevent CHCHD10 accumulation under mitochondrial stress but worsens cell survival. This suggests that CHCHD2 upregulation serves as a protective mechanism under pathological conditions, in line with two recent reports showing that CHCHD2 confers protection to Huntington’s disease models ^62,63^.

Overall, the present study advances our understanding of how loss-of-function mutations in *CHCHD2* contribute to disease pathology, although the mechanisms regulating CHCHD2 levels and assembly remain to be elucidated.

## Materials and Methods

### Ethical statement

All procedures involving animals were conducted following the ethical standards and European, national and institutional guidelines. Protocols were approved by the Stockholm ethical committee and animal work was performed following the guidelines of the Federation of European Laboratory Animal Science Associations (FELASA). For CHCHD2 antibody production, animal experiments were conducted in accordance with the German animal welfare law and performed with permission and adherence to all relevant guidelines and regulations of the district government of Upper Bavaria (Bavaria, Germany; Animal protocol number ROB-55.2Vet-2532.Vet_03-17-68).

### Generation of a monoclonal antibody against CHCHD2

A Wistar rat was immunized with a mixture of 60 µg purified his-tagged mouse CHCHD2 protein in 400 µl PBS, 5 nmol CpG2006 (TIB MOLBIOL, Berlin, Germany), and 400 µl incomplete Freund’s adjuvant. A boost without Freund’s adjuvant was given 12 weeks after primary immunization. Fusion of the myeloma cell line P3X63-Ag8.653 with the rat immune spleen cells was performed 3 days later according to standard procedure ^64^. Hybridoma supernatants were screened 10 day later in a flow cytometry assay (iQue, Intellicyt; Sartorius) for binding to CHCHD2 protein captured on beads (3D-Carboxy, PolyAn, Berlin). Antibody binding was analyzed using ForeCyt software (Sartorius). Positive supernatants were further validated by Western blot on mouse wildtype and ChcHd2 knock-out tissue. Hybridoma cells of clone CCHD 16D9 (rat IgG2a/k) were subcloned twice by limiting dilution to obtain a stable monoclonal cell line.

### Mouse models

Homozygous mice for a *LoxP*-flanked *Chchd2* allele (*Chchd2^loxP/lo^*^xP^) were crossed to heterozygous β-actin-cre to generate whole-body heterozygous *Chchd2* knockout (*Chchd2^+/−^*) mice, which were inter-crossed to obtain homozygous knockout mice (*Chchd2^−^***^/^***^−^*). Animals were housed in a 12-hour light/dark cycle at 21°C and fed with a standard diet *ad libitum*. Analyses of control and knockout mice were performed at different time points. All mice were on the C57BL/6N background.

Snap frozen tissues and mitochondria isolated from i) heart and skeletal muscle tissue-specific *Tefm* knockout mice (*Tefm ^loxP/lo^*^xP^, +/*Ckmm-cre*) at 8 weeks of age ^65^, ii) heart and skeletal muscle tissue-specific *RNAseH1* knockout mice (*RNaseH1 ^loxP/lo^*^xP^, +/*Ckmm-cre*) at 24 weeks of age ^44^ iii) skeletal muscle tissue-specific *Tfam* knockout mice (*Tfam^loxP/lo^*^xP^, +/*Mlc1f-cre*) at 21 week of age ^66^, iv) liver of mice treated with the inhibitor of mitochondrial transcription (IMT) for four weeks ^67^ were obtained from Prof. Nils-Göran Larsson.

### Behavioral tests

Behavioral analyses in control (*Chchd2^+/+^*) and knockout (*Chchd2^−/−^*) mice were performed at 6, 9, 12 and 20 months of age. Distinct testing sessions were conducted for female and male animals. Following an acclimation period of at least 30 min in the ventilated experimental room, each test/training session was performed at the same time during the day (9 am-5 pm).

### Open field and rotarod assays

Animal cohorts were tested for spontaneous locomotion in an open field arena using the ActiMot detection system (TSE systems). Free and uninterrupted movements were recorded for 60 minutes, and spontaneous horizontal/vertical activities and total distance traveled were calculated. Motor coordination was assessed by using a rotarod device (Ugo Basile, Italy). Two training sessions of 90 seconds at the fixed rotation speed of 4 rpm were conducted a day before to the assessment. On the test day, mice were placed on the rod in acceleration mode (4-40 rpm) for 5 minutes and the latency to fall off the rod was recorded. Each animal had five trials with a 5-minute break in between trials.

### Grip-Strength Test and Catwalk

Muscle strength in both the fore- and hindlimbs was tested using the grip strength apparatus (BioSeb, FL, USA). Mice were gently held by the tail base and encouraged to grip a metal grid using their forepaws and hind paws. A horizontal force was applied pulling the mice’s tails. The peak force in grams (g) was quantified using a digital force gauge linked to the metal grid. Each animal was tested three times with a 5-minute break between each measurement. Gait analysis was performed using the CatWalk device (Noldus, Netherlands). Prior to testing, mice underwent a training phase spanning two days to adapt to the test environment. The CatWalk system tracked gait patterns by an enclosed walkway with a glass plate, illuminated using fluorescent light, and equipped with a high-speed color camera. On the testing day, each mouse was stationed at the entrance of the tunnel, allowing unrestricted movement back and forth along the walkway. The protocol mandated the completion of three successful runs within 10 minutes with a duration between 0.5 and 7 seconds and speed variation between 60 and 66 %. Mice unable to meet these criteria were excluded from subsequent analysis.

### High-performance liquid chromatography (HPLC)

Sample preparation and HPLC with electrochemical detection (ECD) were done as previously described (He et al., 2023; Mantas et al., 2020). Catecholamine content of mouse prefrontal cortex, hippocampus, striatum and cerebellum was measured as previously described ^70,71^. In brief, different brain regions were homogenized in ice-cold perchloric acid, after centrifugation, the clear eluents were subjected to HPLC-ECD. Calibration curves were generated using standard solutions to quantify catecholamine content in the different sections of the mouse brain. The separation of analytes was achieved using a reversed-phase C18 column, and detection was performed at specific voltages in an analytical cell, with chromatograms acquired and analyzed to determine the concentration of various neurotransmitters and metabolites. Chemicals and reagents, including DA, HVA, 3-MT, DOPAC, 5-HT, 5-HIAA, DOPA, EPI, NE, MHPG, 70% perchloric acid, 85% phosphoric acid and sodium bisulfite were purchased from Sigma Aldrich.

### Autoradiographic detection of DAT

Fresh frozen sections mounted on Superfrost slides were used for autoradiographic detection of dopamine transporter DAT, as previously described (He et al., 2023).

### Sample preparation for MALDI-MSI

Fresh frozen mouse brains from WT (n=9) and CHCHD2 KO (n=7) were cut at a thickness of 12 μm at prefrontal cortex and striatum levels using a cryostat microtome (Leica CM, Leica Microsystems). Tissue sections were thaw-mounted onto conductive indium tin oxide (ITO) glass slides (Bruker Daltonics) and stored at −80 °C. Sections were desiccated at room temperature for 20 min before spray coating of norharmane matrix solution. The slides were scanned on a flatbed scanner (Epson Perfection V500) after the matrix coating. Norharmane matrix solutions were prepared by dissolving and briefly sonicating the norharmane matrix powder (Sigma Aldrich, N6252) in 80% MeOH (7.5 mg/ml) solution in a glass vial. An automated pneumatic sprayer (HTX-Technologies LLC, Chapel Hill, NC, USA) was used, which was combined with a pump (AKTA FPLC P-905 pump, Cytiva, Uppsala, Sweden) to spray heated matrix solution over the tissue sections. Before the experiments, the pump was kept running at 6 μL/min overnight in between the experiments and 70 μL/min for 2 hours before each experiment using a 50% acetonitrile pushing solvent to ensure a stable flow of the solvent with isocratic pressure. The matrix solution was sprayed using instrumental parameters of a solvent flow rate of 70 μL/min at isocratic pressure, a nitrogen flow of 6 psi, spray temperature of 60 °C, 15 passes (all horizontal), a nozzle head velocity of 1200 mm/min, and track spacing of 2.0 mm.

### MALDI-FTICR-MSI Analysis of Lipids

MALDI-MSI experiments for lipid imaging were performed as previously described ^73^. Briefly, MSI data were collected in dual polarity (both negative and positive ionization modes) on the same tissue sections using a MALDI-FTICR (Solarix XR 7T-2ω, Bruker Daltonics) mass spectrometer equipped with a Smartbeam II 2 kHz laser. The size of the laser was chosen to give a lateral resolution of 100 μm in both polarities with an offset value of 50 μm accompanying the polarity switch. The instrument was tuned for optimal detection of lipid molecules (m/z 200–2000) in both polarities using the quadrature phase detection (QPD) (2ω) mode. The time-of-flight (TOF) values were set at 1.0 ms for negative and 0.8 ms for positive ion mode analysis and the transfer optics frequency was kept at 4 MHz for both polarity analyses. The quadrupole isolation m/z value (Q1 mass) was set at m/z 220.00 for both polarity analyses. Spectra were collected by summing 100 laser shots per pixel in both polarities. Both methods were calibrated externally with red phosphorus over an appropriate mass range. Ion signals of m/z 798.540963 (monoisotopic peak of [PC(34:1)+K]^+^) and m/z 885.549853 (monoisotopic peak of [PI(38:4)-H]^−^) were used for internal calibration for positive and negative polarity analysis, respectively. The data were initially collected in negative polarity and a 5-minute-long dummy experiment in the positive ion mode was performed to ensure the instrument performance after polarity switch. The laser power was optimized at the start of each analysis and then held constant during the MALDI-MSI experiment. Any possible bias due to factors such as matrix degradation or variation in mass spectrometer response was minimized by randomized analysis of the tissue sections.

### MALDI-MSI Data Processing and Statistical Analysis

MSI data were initially visualized in FlexImaging (v.5.0, Bruker Daltonics). For further analysis, data from WT (n=9) and CHCHD2 KO (n=7) were imported to SCiLS Lab (v. 2024a Pro, Bruker Daltonics) and combined. Brain regions were annotated according to the Allen Mouse Brain Atlas ^74^. We utilized the sliding window function of SCiLS Lab to extract root mean square (RMS) normalized average peak area values for 1435 peaks in the negative ion mode and 1229 peaks in positive ion mode separately from each region of interest including the dorsal striatum, ventral striatum and cortex. The combined 2664 peak list in the mass range m/z 400–2000 was combined into an Excel file for multivariate data analysis. GraphPad Prism 10.2.3 (GraphPad Software, La Jolla, California, USA) was used to test data for normality, log_2_-transform data, run multiple t-tests and graph volcano plots. There were no statistically significant results when correcting for multiple comparisons (FDR, Q=5%), so correction was omitted. The resultant peak hits were then visualized in bar graphs by normalizing the non-log transformed RMS peak area values to controls.

### Identification of Lipids

A search of the significant *m/z* values from statistical analysis with a 0.01 *m/z* mass tolerance in LIPID MAPS was conducted for both negative and positive polarities including all the ion types. MALDI-tandem MS (MS/MS) was performed on tissues by collecting spectra from brain regions where the target ion was abundant. The resulting product ions were then compared to product ion spectra of standards in the LIPID MAPS database (Nature Lipidomics Gateway, www.lipidmaps.org) and/or previously published data. In cases where sodium and/or potassium adduct ions of the same lipid species were identified, the brain tissue distribution of the adducts ions and [M+H]^+^ or [M-H]^−^ ions were also compared (Supplementary Table 6) ^37^.

### Cell Lines

Flp-In T-REx HEK293 (HEK293T, RRID: CVCL_U427, female) were grown in DMEM+GlutaMAX culture media (Thermo Fisher, Cat. no. 31966-021) supplemented with 10% fetal bovine serum (Thermo Fisher, Cat. no. 10270-106), 100 U/ml Penicillin / 100 μg/ml Streptomycin (Thermo Fisher, GIBCO, Cat. no. 15140122) and maintained at 37°C and 5% CO_2_. Cells were passaged every 3 to 4 days. Primary mouse embryonic fibroblasts (MEFs) were isolated from embryos at gestational days 13.5 and maintained in culture in the same medium for about 4 passages.

### Assessment of cell growth and viability

Cells were plated in 96-well plates at a density of 1000-2000 cells per well and (10.000 cells) treatments were administered 24 hours after seeding. For growth in galactose, cells were cultured in glucose-free DMEM supplemented with 0.9 g/L galactose, 10% (v/v) FBS, 1 mM sodium pyruvate, 1mM Uridine, 1 x Penicillin/Streptomycin for 96 hours. For cell viability experiments, cells were treated either with 250 nM rotenone for up to 72 hours or with 150 µM actinonin for up to 118 hours.

Cell viability was determined using Cell Counting Kit 8 (Sigma, Cat. no. 96992), which selectively stains viable cells.

### Isolation of mitochondria from mouse tissue

Mice were euthanized by cervical dislocation and several tissues, including brain, heart, skeletal muscle and spinal cord, were collected and washed with ice-cold phosphate-buffered saline (PBS). Mitochondria were isolated using differential centrifugation. Tissues were cut and gently homogenized within in mitochondrial isolation buffer (MIB: 320 mM sucrose, 10 mM Tris–HCl pH 7.4, 1 mM EDTA) supplemented with 1× Complete protease inhibitor cocktail (Roche) and 0,2% of fatty acid-free bovine serum albumin (BSA) using Schuett homgenplus (Schuett Biotec, cat. no. 3.201-011). The tissue homogenate was centrifuged at 1,000 g for 10 minutes at 4°C to remove the cell debris and nuclei. The supernatant containing mitochondria was transferred into a clean, tube and centrifuged at 10000 g for 10 minutes at 4°C to pellet mitochondria. After an additional wash in MIB without BSA, crude mitochondrial pellets were resuspended in a small volume of MIB without BSA, snap-frozen in liquid nitrogen and stored at −80°C.

### Isolation of mitochondria from cultured cells

Cultured cells at 80-90% confluency were scraped from 2-3 150 mm dishes, washed twice in cold PBS and then centrifuged at 800 g for 5 minutes at 4°C. Cell pellet was then resuspended in isolation buffer (CMIB: 20 mM HEPES pH 7.6, 220 mM mannitol, 70 mM sucrose, 1 mM EDTA) supplemented with 1× Complete protease inhibitor cocktail (Roche) and 2 mg/ml of fatty acid-free bovine serum albumin (BSA). To facilitate swelling, the cell suspension was incubated on ice for 15 minutes. Subsequently, homogenization was performed with 20 strokes by using a manual homogenizer. The cell homogenate was centrifuged at 800 g for 5 minutes at 4°C to remove the cell debris. The supernatant was collected into a new tube and centrifuged at 10000 g for 10 minutes at 4°C. The mitochondrial pellet was washed in CMIB without BSA and centrifuged at 10000 g for 10 minutes at 4°C. Mitochondrial extracts were then resuspended in a small volume of CMIB and used for protein quantification. Protein quantification was conducted using a QubitTM Fluorometer (Thermo Fisher, MA, USA).

### Dual COX/SDH enzyme histochemistry

Tissues were frozen in isopentane (15 s) that was previously cooled to −160°C in liquid nitrogen. Tissues were cryosectioned at −18° to −20°C using the Thermo Scientific Microm HM560 cryostat (Thermo Scientific, MA, USA) and were placed onto Polysine microscope slides (VWR, PA, USA, Cat. no. 631-0107). COX/SDH staining was performed as previously described ^75^.

### Biochemical evaluation of respiratory chain function and ATP production

Respiratory chain enzyme activities were measured in isolated mitochondria from different mouse ‘tissues as previously described ^76^.

### Transmission electron microscopy

Small pieces from the myocardium or quadriceps were fixed in 2% glutaraldehyde and 1% paraformaldehyde in phosphate buffer, at room temperature for 30 min, followed by 24 hours at 4°C. Samples were rinsed and subjected to post-fixation in 2% osmium tetroxide in phosphate buffer at 4°C for 2 hours. The samples were then subjected to stepwise ethanol dehydration followed by acetone/LX-112 infiltration and embedded in LX-112 block. Ultrathin sections were prepared using an EM UC7 ultramicrotome (Leica Microsystems). The ultrathin sections were contrasted with uranyl acetate followed by Reynolds lead citrate and examined Hitachi HT7700 microscope (Hitachi, Japan).

### Western blot analysis

Five to twenty micrograms of isolated mitochondria were resuspended in 4x commercial Laemmli buffer protein and run into Bolt^TM^ 4-12% bis-tris Plus protein gels (Thermo Fisher, MA, USA, Cat. no. NW00120BOX) using MES or MOPS running buffer under denaturing conditions. Proteins were transferred onto nitrocellulose membrane using iBlot™ 2 Transfer Stacks system (Thermo Fisher, MA, USA, Cat. no. IB23001). Immunodetection was performed according to standard techniques using enhanced chemiluminescence Immun-Star HRP Luminol/Enhancer (Bio-Rad) and imaging on an ChemiDoc XRS+ system (Biorad). Specific information on antibodies used in this project are provided in Table 1.

**Table 1:**
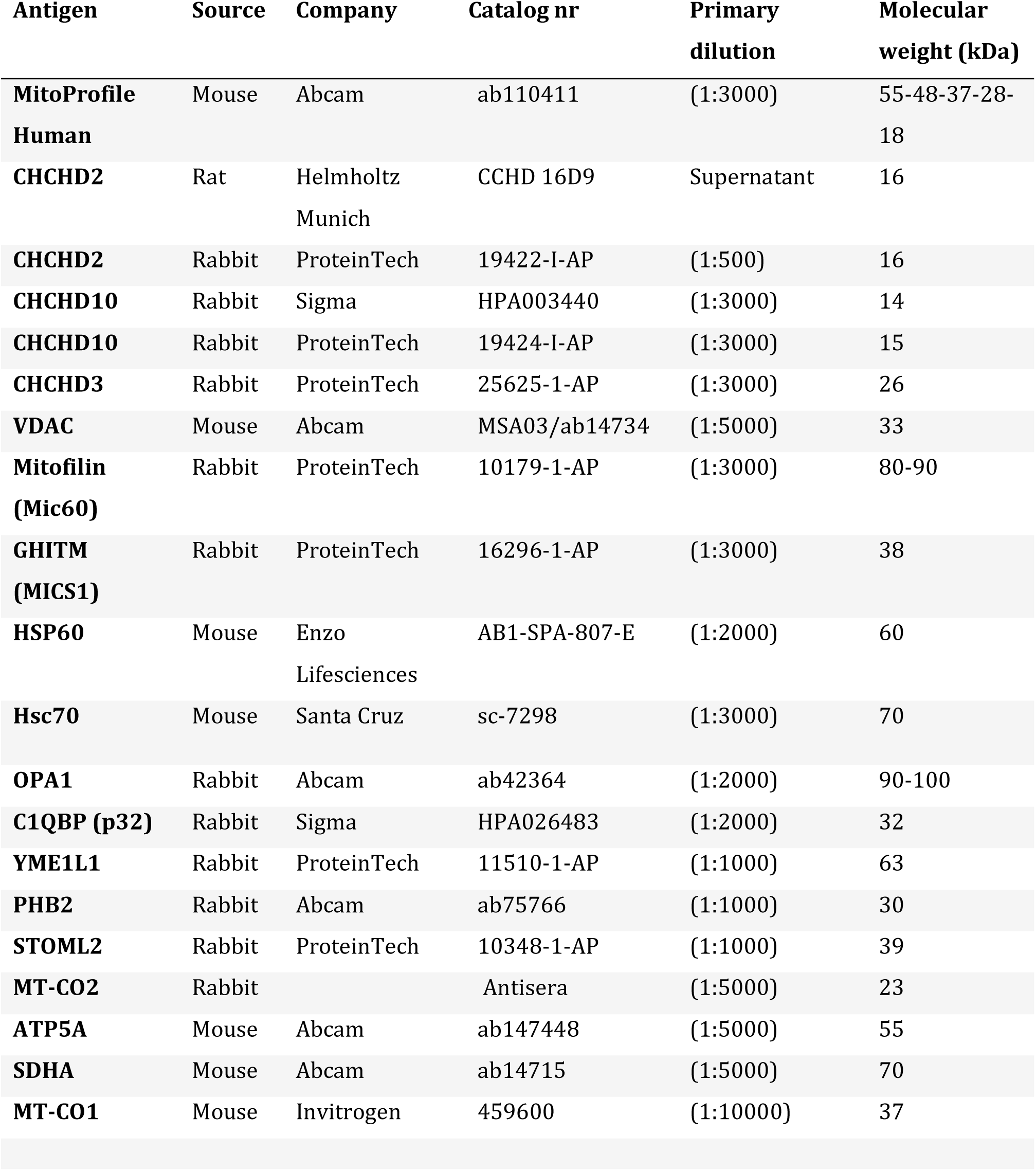
Primary antibody list.

### BN-PAGE and in-gel activity assays

Fifty to hundred micrograms of mitochondria isolated from mouse tissue (liver, skeletal muscle, heart, brain) were lysed in solubilization buffer [20 mM Tris–HCl (pH 7.4), 0.1 mM EDTA, 50 mM NaCl and 10% (v/v) glycerol containing either 1% (w/v) DDM or Digitonin 1% and mixed with loading dye (5% Coomassie Brilliant Blue G-250, 150 mM Bis-Tris and 500 mM 6-aminocaproic acid pH 7.0. Blue native–polyacrylamide gel electrophoresis (BN-PAGE) samples were resolved on 4–16% gradient gels (Thermo Fisher, Cat. no. BN1002BO). The inner chamber was filled with Blue Cathode Buffer [[1x NativePAGE Running buffer (20X), Cat. no. BN2001], [1x NativePAGE™ Cathode Buffer Additive (20X) Catalog number: BN2002], H2O] and the outer chamber with Anode Buffer [1x NativePAGE Running buffer (20X), Cat.no. BN2001, 500 mM Bis-Tris /HCl pH 7.0, 10% digitonin or 10% DDM] at 4°C. BN gels were further subjected to an in-gel catalytic activity assay to measure Complexes I and IV. For complex I activity, the BN-PAGE gels were incubated in 2mM Tris-HCl pH7.4, 0.1 mg/ml NADH (Roche) and 2.5 mg/ml iodonitrozolium for 5-30 minutes. For complex IV activity, BN-PAGE gels were incubated in 0,05 mM phosphate buffer pH7.4, 25 mg 3.3’-diamidobenzidine tetrahydrochloride (DAB), 50 mg cytochrome c, 3,75 g Sucrose and 1 mg Catalase for 1 hour. All steps were carried out at room temperature. Coomassie Imperial Blue staining served as loading control.

### Two-Dimensional Blue Native/SDS Gel Electrophoresis analysis

Strips of the first dimension BN-PAGE gel were incubated for 30 minutes in 1% SDS and 1% β-mercaptoethanol, and then loaded into onto a 1D precast 12% NuPAGETM, Bis-Tris, 1.0 mm, Mini Protein Gel (Thermo Fisher, Cat. no. NP0346BOX) to separate the proteins in the second dimension. The experiment proceeded following the Western blot protocol. Subunits of complex II, IV and V were used to map the molecular weight of different protein complexes.

### Quantification of mtDNA copy number

Genomic DNA was isolated from snap-frozen heart and skeletal muscle (quadriceps) using the DNeasy Blood and Tissue Kit (Qiagen), according to the manufacturer’s instructions. Quantification of mtDNA copy number was performed in triplicates using five ng of DNA using TaqMan Universal Master Mix II and TaqMan probes from Life Technologies. The mtDNA levels were assessed using probes against the mitochondrial genes (ND1 and ND4) and nuclear 18S rRNA gene was used as a loading control. (TaqMan Probes are listed in Table)

### Immunoprecipitation (IP) and mass spectrometry analysis

Immunoprecipitation experiments were performed using various antibodies coupled with the Dynabeads Protein G system (Thermo Fisher 10004D), according to manufacturer’s instructions. Briefly, for 1 mg of mitochondria isolated from HEK 293T cells, 50μL of DynaBeads were employed and an additional 12μL were used for the preclearing process. For optimal antibody binding to the beads, 1 μg of the chosen antibody was incubated with Dynabeads Protein for 1 hour at 4°C. Mitochondria extracts were prepared in lysis buffer (10 mM Tris–HCl pH 7.5, 150 mM NaCl), supplemented with 1% Digitonin and protease inhibitor (PI). After a centrifugation step at 14000 g at 4°C for 10 minutes, the supernatant containing the mitochondrial lysates fractions was pre-cleared for 30 minutes with non-coated Dynabeads Protein G to reduce non-specific protein binding to the beads. The immunoprecipitation reaction between precleared lysate and Dynabeads Protein G bound with indicated antibodies was performed for 2 hours at 4°C. After extensive washes, one-quarter of the samples were eluted with 1% SDS for 10 minutes at room temperature and further employed for Western blot analysis. The remaining beads were analyzed by mass spectrometry. Peptide MS/MS spectra of trypsin-digested proteins were detected. Collected raw data was preprocessed by using MaxQuant software without initial normalization to allow the detection of low-abundant proteins ^77^. Reviewed Swiss-Prot Proteomes from experimental organisms in FASTQ-format were used as reference proteomes ^78^. Perseus software was employed for downstream analysis of collected mass spectrometry data and detected proteins were filtered for mitochondrial subcellular localization using MitoCarta3.0 ^79^. For visualization of volcano plots OmicLoupe (Version v1.1.7) was employed ^80^.

### Complexome profiling

Isolated mitochondria from quadriceps were subjected to complexome profiling proteomics after resolving of protein complexes by 6-16.5% BN-PAGE as extensively described in ^81^.

In brief, the BN-PAGE gel lanes were segmented into 64 equal-sized bands and each band was sectioned into 1 mm^3^ pieces. These fragments were then processed in an OASIS® HLB μElution Plate, where they underwent de-staining, reduction, alkylation, tryptic digestion, and desalting. The resulting peptides were suspended in 0.1% formic acid and analyzed via mass spectrometry using an Exploris 480 mass spectrometer coupled with an EASY-nLC 1200 UHPLC system. Peptides were separated on a reversed-phase C18 column (Auoroa Frontiers, 15cm, 1.7 µm, IonOpticks) over a total 30 min gradient run. Mass spectra were acquired in a Top15 data-dependent manner, with fragmentation achieved through HCD. The RF levels was set to 45 and the capillary temperature was set to 275°C. MS1 spectra were acquired at a resolution of 120K using an AGC (Automatic gain control) target of 1e6. The MS2 spectra were acquired using a resolution of 15K, a maximum injection time of 23 ms, and an AGC target of 5e5. Raw data were analyzed using MaxQuant version 2.4.2.0, searching against the Uniprot mouse reference proteome (one fast per gene) containing 21,984 protein sequences (downloaded June 2020). The match between runs algorithm was enabled using default settings. Protein quantification was conducted using iBAO intensity. The data were scaled row-wise between 0-1 for representation in a heatmap or line diagram using the Instant Clue Software ^82^.

### RNA extraction and RT-qPCR

Total RNA was isolated from snap-frozen mouse tissues (heart, skeletal muscle and liver) using TRIzol (Thermo Fisher Scientific) and quantified by Nanodrop spectrophotometer. Reverse transcription was performed using the High-Capacity cDNA Reverse Transcription Kit (Applied Biosystems, Life Technologies). qPCR was performed on a QuantStudio 6 (Thermo Fisher Scientific) using the TaqMan Universal Master Mix II with the following TaqMan probes (listed in Table 2) from Life Technologies. β-Actin was used as a loading control.

**Table 2.** List of TaqMan probes.

| **GENE** | **TAQMAN PROBE ID** |
| --- | --- |
| *Chchd2* | Mm01621550_s1 |
| *Chchd10* | Mm07302070_g1 |
| *Mthfd2* | Mm00485276_m1 |
| *B- Actin* | Mm01205647_g1 |
| *Mt-Co1* | Mm04225243_g1 |
| *Cytb* | Mm04225271_g1 |
| *Nd1* | Mm04225274_s1 |
| *Nd4* | Mm04225294_s1 |
| *Atf4* | Mm00515325_g1 |
| *Atf5* | Mm04179654_m1 |
| *Chop Or Ddit3* | Mm01135937_g1 |
| *Fgf21* | Mm00840165_g1 |
| *18s RRNA* | Mm03928990_g1 |

**Table 3.** List of interactors identified in endogenous CHCHD2 from HEK293T cells.

| *Gene name* | *-Log(P-value)* | *lof2FC* | *Protein names* |
| --- | --- | --- | --- |
| *C1QBP* | 6,075158655 | 50,33133443 | Complement component 1 Q subcomponent-binding protein, mitochondrial |
| *CHCHD2* | 4,924605406 | 46,57842827 | Coiled-coil-helix-coiled-coil-helix domain-containing protein 2, mitochondrial |
| *WARS2* | 11,28814824 | 21,0234251 | Tryptophan--tRNA ligase, mitochondrial |
| *VDAC1* | 13,49487394 | 20,67803733 | Voltage-dependent anion-selective channel protein 1 |
| *POLDIP2* | 12,47333063 | 19,60814031 | Polymerase delta-interacting protein 2 |
| *GHITM* | 2,834959277 | 19,44199308 | Growth hormone-inducible transmembrane protein |
| *PHB2* | 3,531659873 | 19,00003929 | Prohibitin-2 |
| *STOML2* | 2,836382307 | 18,47478898 | Stomatin-like protein 2, mitochondrial |
| *MT-CO2* | 2,839913569 | 18,36949666 | Cytochrome c oxidase subunit 2 |
| *SLC25A5* | 2,830006645 | 18,11589368 | ADP/ATP translocase 2;ADP/ATP translocase 2, N-terminally processed |
| *SLC25A3* | 2,833964049 | 17,96081797 | Phosphate carrier protein, mitochondrial |
| *SFXN1* | 2,837556211 | 17,75948079 | Sideroflexin-1 |
| *CHCHD10* | 2,791752512 | 17,32434845 | Coiled-coil-helix-coiled-coil-helix domain-containing protein 10, mitochondrial |
| *PHB* | 2,264291177 | 14,97139657 | Prohibitin |

**Table 4.** List of interactors identified in endogenous CHCHD10 from HEK293T cells.

| *Gene names* | *-logP* | *logFC* | *Protein names* |
| --- | --- | --- | --- |
| *CHCHD10* | 13,0391499 | 29,95812511 | Coiled-coil-helix-coiled-coil-helix domain-containing protein 10, mitochondrial |
| *SLC25A11* | 13,00038678 | 22,20192003 | Mitochondrial 2-oxoglutarate/malate carrier protein |
| *CHCHD2* | 12,70984575 | 29,81633472 | Coiled-coil-helix-coiled-coil-helix domain-containing protein 2, mitochondrial |
| *GHITM* | 9,470368995 | 24,75180149 | Growth hormone-inducible transmembrane protein |
| *SLC25A1* | 8,83733878 | 21,1583395 | Tricarboxylate transport protein, mitochondrial |
| *PHB* | 3,042742019 | 1,763472271 | Prohibitin |
| *SLC25A6;*<br>*SLC25A4* | 2,716956338 | 19,78556242 | ADP/ATP translocase 3;ADP/ATP translocase 3, N-terminally processed;ADP/ATP translocase 1 |
| *HSPA9* | 2,653117582 | 1,753324604 | Stress-70 protein, mitochondrial |
| *ATP5A1* | 2,464498305 | 18,51404667 | ATP synthase subunit alpha, mitochondrial |
| *PHB2* | 2,40315313 | 19,76722679 | Prohibitin-2 |
| *VDAC2* | 2,391657039 | 18,34489145 | Voltage-dependent anion-selective channel protein 2 |
| *HSP90AA1* | 2,16463086 | 16,3484334 | Heat shock protein HSP 90-alpha |
| *PGAM5* | 2,008958022 | 17,22167692 | Serine/threonine-protein phosphatase PGAM5, mitochondrial |
| *ATP5C1* | 1,958327291 | 15,98890734 | ATP synthase subunit gamma, mitochondrial |
| *MT-CO2* | 1,949919387 | 16,92953634 | Cytochrome c oxidase subunit 2 |
| *COX4I1* | 1,947517998 | 16,19647694 | Cytochrome c oxidase subunit 4 isoform 1, mitochondrial |
| *SFXN1* | 1,934530549 | 15,73735237 | Sideroflexin-1 |
| *STOML2* | 1,927370711 | 16,13535681 | Stomatin-like protein 2, mitochondrial |
| *PARL* | 1,925565354 | 14,82555056 | Presenilins-associated rhomboid-like protein, mitochondrial;P-beta |
| *P32* | 1,921266933 | 19,99099941 | Complement component 1 Q<br>subcomponent-binding protein,<br>mitochondrial |
| *ATP5B* | 1,853779802 | 15,60795879 | ATP synthase subunit beta, mitochondrial |
| *YME1L1* | 1,577678431 | 15,16261683 | ATP-dependent zinc metalloprotease<br>YME1L1 |

**Table 5:** List of common interactors between CHCHD2 and CHCHD10.

| *Gene name* | *CHCHD10: log2FC* | *CHCHD2: log2FC* |
| --- | --- | --- |
| **CHCHD10** | 20,4757069 | 17,3243484 |
| **CHCHD2** | 19,9869816 | 46,5784283 |
| **GHITM** | 17,2734874 | 19,4419931 |
| **PHB2** | 13,8306508 | 19,0000393 |
| **STOML2** | 13,2640279 | 18,474789 |
| **ATP5C1** | 13,1421083 | 8,74178092 |
| **SFXN1** | 12,5898819 | 17,7594808 |
| **MT-CO2** | 10,6908516 | 18,3694967 |
| **ATP5B** | 9,81515361 | 12,305958 |
| **C1QBP** | 9,76848689 | 50,3313344 |
| **ATP5A1** | 9,63199343 | 8,08011036 |
| **YME1L1** | 9,50194866 | 15,4506259 |
| **MRPL34** | 8,96722505 | 4,72670574 |
| **CYC1** | 2,56963337 | 13,0757624 |
| **ATAD3A** | 2,49557571 | -4,73028011 |
| **SLC25A5** | 1,77081964 | 18,1158937 |
| **HSPA9** | 1,49343104 | 0,15775312 |
| **PHB** | 1,43103698 | 14,9713966 |
| **MRPS12** | 0,65431851 | 0,06124662 |
| **MRPL14** | -0,00709872 | 0,64255511 |

### Statistical analysis

GraphPad Prism v10 software (version 10.0.3) was used for statistical analyses and graph plotting. All data are presented as means ± SEM and statistical comparisons were performed using unpaired t-test. The number of animals or biological replicates (n) used for each experiment is indicated in the figure legends. The statistical analysis of MS spectrometry data comparing IP pull-down samples and controls was carried out using both-sided t-test. Values of p < 0.05 were considered statistically significant.

## Acknowledgments

We would like to thank Prof. Nils-Göran Larsson, Karolinska Institutet, who kindly supplied tissues and isolated mitochondria from different mouse models. We thank Florian Rosenberger, Max Planck Institute of Biochemistry, for excellent technical help in proteomic analysis. We appreciate the support from Proteomics Biomedicum core facility (A. Vegvari) and Electron microscopy core facility (L. Haag), Karolinska Institutet, and Science for Life Laboratory, Spatial Mass Spectrometry, Uppsala University, Uppsala, Sweden. We thank Dominique Diehl for expert technical assistance, Max Planck Institute for Biology of Ageing. Figures have been created using BioRender.com.

This study was supported by grants to R.Fi. from Vetenskapsrå det (2022-01477), Hjärnfonden, Loo och Hans Ostermans stiftelse, Å hlén-stiftelsen, KI Research Foundation Grants, StratNeuro and Hedlunds stiftelse, Karolinska Institutet KID Funding Program for doctoral students. P.E.A was supported from Vetenskapsrå det (2021-03293) and Science for Life Laboratory. P.S. was supported by the Stockholm City Council and Knut och Alice Wallenbergs Stiftelse.

## Author contributions

Methodology: J.G., P.P., X.Z., S.W., M.M., D.M., H.N., I.K., R.G.V., D.R.G., D.A., and R.Fe. Analyses: J.G., P.P., X.Z., S.W., M.M., D.M., H.N., I.K., and R.Fi. Writing Original Draft: J.G., P.P., and R.Fi. Review & Editing: all authors. Visualization: J.G., P.P., and R.Fi. Supervision and Funding Acquisition: P.E.A., T.L., P.S., and R.Fi.

## Data and materials availability

All data needed to evaluate the conclusions in the paper are present in the paper and/or the Supplementary Materials.

## Figure Legends

**Fig. S1. Generation of *Chchd2* Knockout mice. A.** Targeting strategy for disruption of the *Chchd2* gene. LoxP sites flanking exons 2 and 3 of the *Chchd2* gene together with neomycin selection marker (Neo R) were inserted into the mouse genome by homologous recombination. The Neo R cassette was excised by mating with flp-recombinase mice to obtain heterozygous floxed *Chchd2* mice (*Chchd2 ^+/loxP^*). Whole-body *Chchd2* knockout mice were obtained by crossing *Chchd2 ^+/loxP^* with Beta-actin cre-recombinase mice. **B.** Mendelian distribution of offspring obtained by crossing heterozygous knockout *Chchd2* (*Chchd2^+/-^)* males and females. n = 212; *Chchd2^−/−^* n = 53; *Chchd2^+/-^* n = 104; *Chchd2^+/+^* n =55. **C.** Western blot of the CHCHD2 protein levels in mitochondrial extracts from different tissues of one-month-old *Chchd2^+/+^* and *Chchd ^−/−^* mice. VDAC was used as loading control. **D.** Quadriceps (QW) and heart (HW) weight to BW (g/g) ratios in males at 12 months of age. Data are represented as means ± SEM; n>18, ns not significant. **E.** BW (g) of female *Chchd2^−/−^* and *Chchd2^+/+^* mice at the age of 20 months. Data are represented as means ± SEM; n≥ 5, ns not significant.

**Fig. S2. Spontaneous locomotion is not affected by the loss of CHCHD2 in both male and female mice.** Spontaneous locomotor activity measured as distance and rearing, in open field arena over 60 minutes in **A.** *Chchd2^−/−^*and *Chchd2^+/+^* male mice at the age of 6, 9 and 12 months. Data are represented as means ± SEM; n ≥ 16, ns not significant. **B.** in *Chchd2^−/−^* and *Chchd2^+/+^* female mice at the age of 20 months. Data are represented as means ± SEM; *n* = 5–7; ns not significant. **C.** Motor coordination measured as latency to fall (in s) from the rod and **D.** Muscular strength (in g) measured by grip strength test in 20-month-old female mice. Data are represented as means ± SEM *n* = 5–7; ns not significant.

**Fig. S3. Analysis of monoamine neurotransmitters in the brain of male and female mice lacking CHCHD2. A**. Hippocampal levels (ng/mg) of 3-Methoxy-4-hydroxyphenylglycol (MHPG), epinephrine (EPI) and 5-Hydroxyindoleacetic acid (5-HIAA) **B.** Cerebellar levels (ng/mg) of NE in *Chchd2^−/−^* and *Chchd2^+/+^* male mice at the age of 12 months. Data are represented as means± SEM; n ≥ 7, *p < 0.05, ns not significant. **C.** Heatmaps showing the levels (ng/mg) of DA, NE, 5-HT and their metabolites in the prefrontal cortex, hippocampus, striatum and cerebellum of *Chchd2^−/−^*and *Chchd2^+/+^* female mice at the age of 20 months. **D.** DA turnover ratios (DOPAC/DA, 3-MT/DA and HVA/DA) in the prefrontal cortex of *Chchd2^−/−^*and *Chchd2^+/+^* female mice at the age of 20 months. Data are represented as means ± SEM *n* = 5–7; *p < 0.05, ns not significant.

**Fig. S4. CHCHD2 loss does not compromise OXPHOS nor promote integrated stress response.** Western blot analysis of steady-state levels of **A.** OXPHOS subunits in mitochondria isolated from different tissues of *Chchd2^−/−^* and *Chchd2^+/+^* male mice at the age of 12 months. VDAC was used as loading control. **B.** Complex IV and V in-gel activities performed on mitochondria isolated from different tissues of *Chchd2^−/−^* and *Chchd2^+/+^* male mice at the age of 12 months. BN-PAGE stained with Coomassie or incubated with substrates for detecting the in-gel activity of the indicated OXPHOS complexes. **C.** Cytochrome c oxidase and succinate dehydrogenase (COX/SDH) double-labelling enzyme histochemistry of colon in *Chchd2^−/−^* and *Chchd2^+/+^* mice at the age of 12 months (Scale bar: 100 μm). COX-deficient colonic crypts (in blue) are indicated by the yellow arrows. **D.** *Atf4, Atf5, Chop* and *Fgf21* transcript levels measured by qPCR on RNA isolated from heart and skeletal muscle (quadriceps) in mice at 12 months of age. Data are represented as means ± SEM; *n*=5; ns not significant.

**Fig. S5. Mild alterations in lipid homeostasis in male mice lacking CHCHD2. A.** Relative quantification of affected lipids in different brain regions of *Chchd2^−/−^* male mice at the age of 12 months. Data are represented as means ± SEN; *n* ≥ 7; *p< 0.05, **P < 0.01.

**Fig. S6. CHCHD2 and CHCHD10 are not MICOS subunits.** 2D-PAGE of mitochondria isolated **A.** from skeletal muscle of *Chchd2^−/−^* and *Chchd2^+/+^* male mice at the age of 12 months. Mitochondria were solubilized using 1% (w/v) digitonin. **B.** from HEK293T cells and solubilized using either with 1% (w/v) DDM or digitonin. **C.** from MEFs and solubilized using 1% (w/v) DDM. The position of SDHA, ATP5, MT-COI, and CHCHD3 corresponds to the size of their different protein complexes (in kDa). **D.** Mitochondria isolated from skeletal muscle of *Chchd2^−/−^* and *Chchd2^+/+^* male mice at the age of 12 months (n=3-4) were solubilized using 1% (w/v) digitonin and resolved on self-made 3%– 13% gels. **E.** Migration profiles of CHCHD10 obtained from complexome profiling analysis of mitochondria isolated from skeletal muscle of *Chchd2^−/−^* mice at the age of 12 months. n=3. **F.** Heatmaps (upper panel) and migration profiles (lower panel) of the MICOS complex subunits obtained by complexome profiling performed on mitochondria isolated from skeletal muscle of *Chchd2^−/−^* and *Chchd2^+/+^* male mice at the age of 12 months. n=3. **G-H.** Volcano plots of the hits identified by mass spectrometry analysis on **G.** CHCHD2 and **H**. CHCHD10 pull-down experiments. **I.** 2D-PAGE electrophoresis analyses of mitochondria isolated from heart of *Chchd2^+/+^* male mice at the age of 12 months. Mitochondria were solubilized using 1% (w/v) DDM. The position of SDHA, ATP5, MT-CO1, and CHCHD3 (indicated by a red arrow) corresponds to the size of their different protein complexes (in kDa). P32 is indicated by blue arrows.

**Fig. S7. CHCHD2 and CHCHD10 form a larger and more abundant complex in response to mitochondrial dysfunction. A.** Western blot analysis of OXPHOS subunits levels in mitochondria isolated from mouse models with impaired mitochondria. VDAC and Coomassie staining were used as loading controls. **B.** Complex IV and V in-gel activities performed on mitochondria isolated from heart of *Tefm* knockout mice at 8 weeks of age. **C.** *Mthfd2* mRNA levels measured by qPCR from RNA isolated from mouse tissues with mitochondrial dysfunction. Data are represented as mean ± SEM. n= 5 mice for each genotype, *p < 0.05, **p < 0.01, ***p< 0.001. **D.** Western Blot analysis of CHCHCD2 and CHCHD10 steady-state levels in mitochondria isolated from heart of *Tefm* knockout mice at 8 weeks of age. **E.** *Chchd2* and *Chchd10* mRNA levels measured by qPCR from RNA isolated from heart of *Tefm* knockout mice at 8 weeks of age. Data are represented as mean ± SEM. n= 5 mice for each genotype, *p < 0.05. **F.** 2D-PAGE of mitochondria isolated from skeletal muscle of from heart of *Tefm* knockout mice at 8 weeks of age. Mitochondria were solubilized using 1% (w/v) DDM. The position of SDHA, ATP5, MT-CO1, and CHCHD3 (indicated by a red arrow) corresponds to the size of their different protein complexes (in kDa). P32 is indicated by blue arrows.

**Fig. S8. CHCHD2 and CHCHD10 accumulate in MEFs treated with rotenone or actinonin. A-B.** Representative Western Blot analysis of CHCHCD2 and CHCHD10 steady-state levels in total extract from primary *Chchd2^+/+^* MEFs treated with 250 nM rotenone for 24, 48 and 72 hours and 150 nM actinonin for 24, 72 and 118 hours. HSC70 and Coomassie were used as loading control.

---

**Supplementary Table 6.**
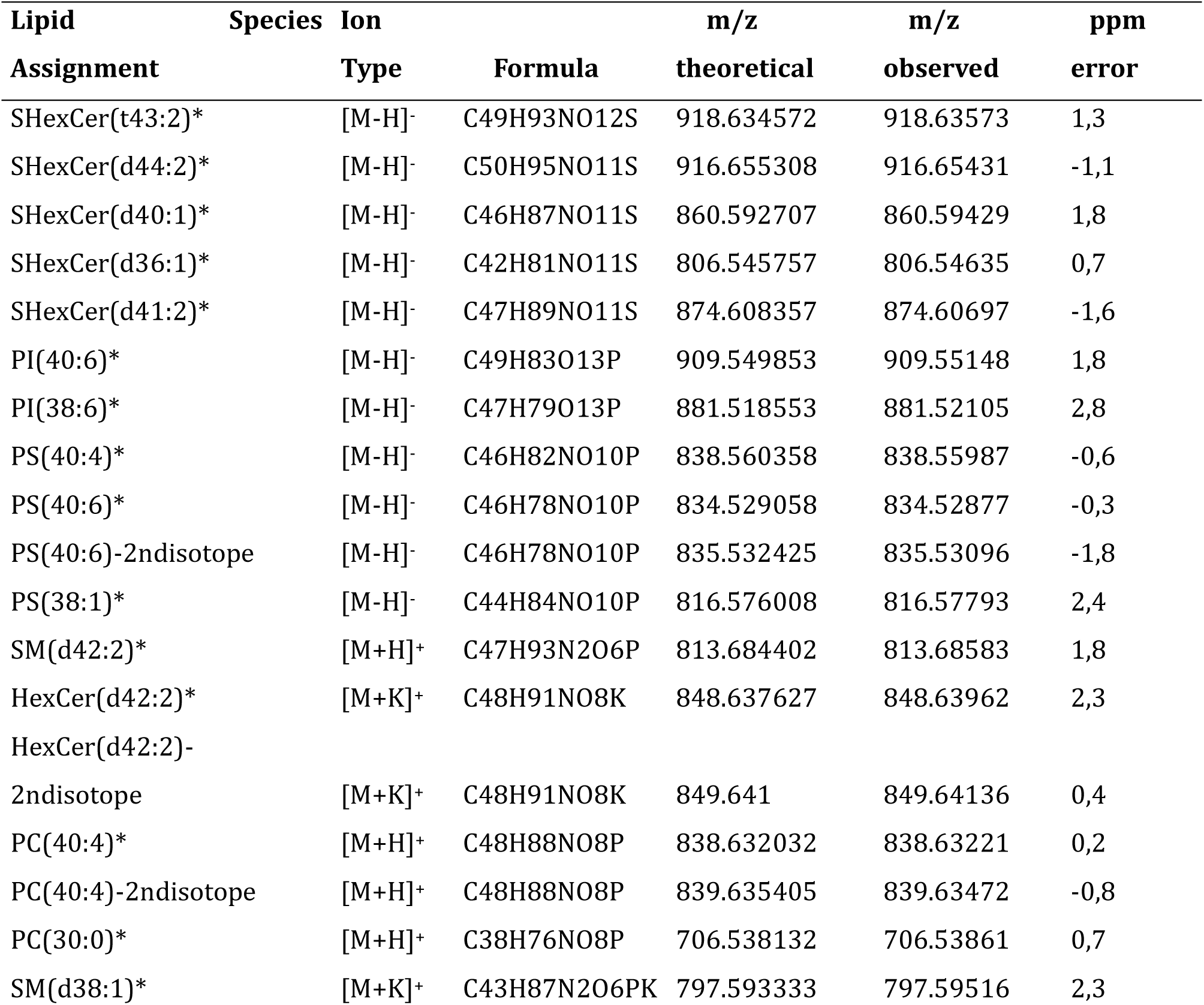
List of assigned lipid species with high mass accuracy obtained from MALDI-FTICR-MSI data. Lipid species marked with an asterisk (*) were identified using mass accuracy, MS/MS and by comparing the observed distributions of different adducts of the same molecule across the coronal brain tissue sections.

## References

1. Hung, V. et al. Proteomic Mapping of the Human Mitochondrial Intermembrane Space in Live Cells via Ratiometric APEX Tagging. Mol Cell 55, 332–341 (2014).

2. Habich, M., Salscheider, S. L. & Riemer, J. Cysteine residues in mitochondrial intermembrane space proteins: more than just import. Br J Pharmacol 176, 514– 531 (2019).

3. Zarges, C. & Riemer, J. Oxidative protein folding in the intermembrane space of human mitochondria. FEBS Open Bio (2024) doi:10.1002/2211-5463.13839.

4. Herrmann, J. M. & Riemer, J. The Intermembrane Space of Mitochondria. Antioxid Redox Signal 13, 1341–1358 (2010).

5. Herrmann, J. M. & Bykov, Y. Protein translocation in mitochondria: Sorting out the Toms, Tims, Pams, Sams and Mia. FEBS Lett 597, 1553–1554 (2023).

6. Modjtahedi, N., Tokatlidis, K., Dessen, P. & Kroemer, G. Mitochondrial Proteins Containing Coiled-Coil-Helix-Coiled-Coil-Helix (CHCH) Domains in Health and Disease. Trends Biochem Sci 41, 245–260 (2016).

7. Rampelt, H., Zerbes, R. M., van der Laan, M. & Pfanner, N. Role of the mitochondrial contact site and cristae organizing system in membrane architecture and dynamics. Biochimica et Biophysica Acta (BBA) - Molecular Cell Research 1864, 737–746 (2017).

8. Funayama, M. et al. CHCHD2 mutations in autosomal dominant late-onset Parkinson’s disease: a genome-wide linkage and sequencing study. Lancet Neurol 14, 274–282 (2015).

9. Yang, N. et al. Systematically analyzing rare variants of autosomal-dominant genes for sporadic Parkinson’s disease in a Chinese cohort. Neurobiol Aging 76, 215.e1–215.e7 (2019).

10. Jansen, I. E. et al. CHCHD2 and Parkinson’s disease. Lancet Neurol 14, 678–679 (2015).

11. Ogaki, K. et al. Mitochondrial targeting sequence variants of the *CHCHD2* gene are a risk for Lewy body disorders. Neurology 85, 2016–2025 (2015).

12. Lee, R. G. et al. Early-onset Parkinson disease caused by a mutation in CHCHD2 and mitochondrial dysfunction. Neurol Genet 4, e276 (2018).

13. Kee, T. R. et al. Mitochondrial CHCHD2: Disease-Associated Mutations, Physiological Functions, and Current Animal Models. Front Aging Neurosci 13, (2021).

14. Shammas, M. K., Huang, T.-H. & Narendra, D. P. CHCHD2 and CHCHD10-related neurodegeneration: molecular pathogenesis and the path to precision therapy. Biochem Soc Trans 51, 797–809 (2023).

15. Jiang, Z. et al. Production of a human iPSC line from an early-onset Parkinson’s disease patient with a novel CHCHD2 gene truncated mutation. Stem Cell Res 64, 102881 (2022).

16. Leventoux, N. et al. Aberrant CHCHD2-associated mitochondriopathy in Kii ALS/PDC astrocytes. Acta Neuropathol 147, 84 (2024).

17. Cavallaro, G. Genome-wide analysis of eukaryotic twin CX9C proteins. Mol Biosyst 6, 2459 (2010).

18. Straub, I. R. et al. Loss of CHCHD10-CHCHD2 complexes required for respiration underlies the pathogenicity of a CHCHD10 mutation in ALS. Hum Mol Genet 27, 178–189 (2018).

19. Burstein, S. R. et al. In vitro and in vivo studies of the ALS-FTLD protein CHCHD10 reveal novel mitochondrial topology and protein interactions. Hum Mol Genet 27, 160–177 (2018).

20. Bannwarth, S. et al. A mitochondrial origin for frontotemporal dementia and amyotrophic lateral sclerosis through CHCHD10 involvement. Brain 137, 2329– 2345 (2014).

21. Johnson, J. O. et al. Mutations in the CHCHD10 gene are a common cause of familial amyotrophic lateral sclerosis. Brain 137, e311–e311 (2014).

22. Ajroud-Driss, S. et al. Mutation in the novel nuclear-encoded mitochondrial protein CHCHD10 in a family with autosomal dominant mitochondrial myopathy. Neurogenetics 16, 1–9 (2015).

23. Penttilä, S., et al. *CHCHD10* mutations and motor neuron disease: the distribution in Finnish patients. J Neurol Neurosurg Psychiatry 88, 272–277 (2017).

24. Auranen, M., et al. *CHCHD10* variant p.(Gly66Val) causes axonal Charcot-Marie-Tooth disease. Neurol Genet 1, (2015).

25. Bannwarth, S. et al. A mitochondrial origin for frontotemporal dementia and amyotrophic lateral sclerosis through CHCHD10 involvement. Brain 137, 2329– 2345 (2014).

26. Genin, E. C. et al. <scp>>CHCHD</scp> 10 mutations promote loss of mitochondrial cristae junctions with impaired mitochondrial genome maintenance and inhibition of apoptosis. EMBO Mol Med 8, 58–72 (2016).

27. Genin, E. C. et al. CHCHD10 and SLP2 control the stability of the PHB complex: a key factor for motor neuron viability. Brain 145, 3415–3430 (2022).

28. Meng, H. et al. Loss of Parkinsons disease-associated protein CHCHD2 affects mitochondrial crista structure and destabilizes cytochrome c. Nat Commun 8, 1– 18 (2017).

29. Nguyen, M. K. et al. Mouse midbrain dopaminergic neurons survive loss of the PD-associated mitochondrial protein CHCHD2. Hum Mol Genet (2021) doi:10.1093/hmg/ddab329.

30. Sato, S. et al. Homeostatic p62 levels and inclusion body formation in CHCHD2 knockout mice. Hum Mol Genet 30, 443–453 (2021).

31. Baines, H. L. et al. Similar patterns of clonally expanded somatic mtDNA mutations in the colon of heterozygous mtDNA mutator mice and ageing humans. Mech Ageing Dev 139, 22–30 (2014).

32. Shammas, M. K. et al. OMA1 mediates local and global stress responses against protein misfolding in CHCHD10 mitochondrial myopathy. Journal of Clinical Investigation 132, (2022).

33. Kühl, I., et al. Transcriptomic and proteomic landscape of mitochondrial dysfunction reveals secondary coenzyme Q deficiency in mammals. Elife 6, (2017).

34. Liu, Y.-T. et al. Loss of CHCHD2 and CHCHD10 activates OMA1 peptidase to disrupt mitochondrial cristae phenocopying patient mutations. Hum Mol Genet 29, 1547– 1567 (2020).

35. Sastry, P. S. Lipids of nervous tissue: Composition and metabolism. Prog Lipid Res 24, 69–176 (1985).

36. He, Y. et al. Prosaposin maintains lipid homeostasis in dopamine neurons and counteracts experimental parkinsonism in rodents. Nat Commun 14, 5804 (2023).

37. Kaya, I. et al. Spatial lipidomics reveals brain region-specific changes of sulfatides in an experimental MPTP Parkinson’s disease primate model. NPJ Parkinsons Dis 9, 118 (2023).

38. Kaya, I. et al. Spatial lipidomics reveals brain region-specific changes of sulfatides in an experimental MPTP Parkinson’s disease primate model. NPJ Parkinsons Dis 9, 118 (2023).

39. Wai, T. et al. The membrane scaffold SLP2 anchors a proteolytic hub in mitochondria containing PARL and the *i*-AAA protease YME1L. EMBO Rep 17, 1844–1856 (2016).

40. Ren, Y.-L. et al. Loss of CHCHD2 Stability Coordinates with C1QBP/CHCHD2/CHCHD10 Complex Impairment to Mediate PD-Linked Mitochondrial Dysfunction. Mol Neurobiol (2024) doi:10.1007/s12035-024-04090-y.

41. Huang, X. et al. CHCHD2 accumulates in distressed mitochondria and facilitates oligomerization of CHCHD10. Hum Mol Genet (2018) doi:10.1093/hmg/ddy270.

42. Jiang, S. et al. <scp>TEFM</scp> regulates both transcription elongation and <scp>RNA</scp> processing in mitochondria. EMBO Rep 20, (2019).

43. Stroud, D. A. et al. Accessory subunits are integral for assembly and function of human mitochondrial complex I. Nature 538, 123–126 (2016).

44. Misic, J. et al. Mammalian RNase H1 directs RNA primer formation for mtDNA replication initiation and is also necessary for mtDNA replication completion. Nucleic Acids Res 50, 8749–8766 (2022).

45. Wredenberg, A. et al. Increased mitochondrial mass in mitochondrial myopathy mice. Proceedings of the National Academy of Sciences 99, 15066–15071 (2002).

46. Jiang, S. et al. Inhibition of mammalian mtDNA transcription acts paradoxically to reverse diet-induced hepatosteatosis and obesity. Nat Metab 6, 1024–1035 (2024).

47. Khan, N. A. et al. mTORC1 Regulates Mitochondrial Integrated Stress Response and Mitochondrial Myopathy Progression. Cell Metab 26, 419–428.e5 (2017).

48. Tiklová, >K., et al. Single-cell RNA sequencing reveals midbrain dopamine neuron diversity emerging during mouse brain development. Nat Commun 10, 581 (2019).

49. Baughman, J. M. et al. A Computational Screen for Regulators of Oxidative Phosphorylation Implicates SLIRP in Mitochondrial RNA Homeostasis. PLoS Genet 5, e1000590 (2009).

50. Aras, S. et al. MNRR1 (formerly CHCHD2) is a bi-organellar regulator of mitochondrial metabolism. Mitochondrion 20, 43–51 (2015).

51. Longen, S. et al. Systematic Analysis of the Twin Cx9C Protein Family. J Mol Biol 393, 356–368 (2009).

52. Petel Légaré, V., et al. Loss of mitochondrial Chchd10 or Chchd2 in zebrafish leads to an ALS-like phenotype and Complex I deficiency independent of the mitochondrial integrated stress response. Dev Neurobiol 83, 54–69 (2023).

53. Lumibao, J. et al. CHCHD2 mediates glioblastoma cell proliferation, mitochondrial metabolism, hypoxia-induced invasion and therapeutic resistance. Int J Oncol 63, 117 (2023).

54. Ruan, Y. et al. CHCHD2 and CHCHD10 regulate mitochondrial dynamics and integrated stress response. Cell Death Dis 13, 156 (2022).

55. Liu, W. et al. Chchd2 regulates mitochondrial morphology by modulating the levels of Opa1. Cell Death Differ 27, 2014–2029 (2020).

56. Ahola, S. et al. Opa1 processing is dispensable in mouse development but is protective in mitochondrial cardiomyopathy. Sci Adv 10, (2024).

57. Southwell, N. et al. High fat diet ameliorates mitochondrial cardiomyopathy in CHCHD10 mutant mice. EMBO Mol Med 16, 1352–1378 (2024).

58. Potting, C. et al. TRIAP1/PRELI Complexes Prevent Apoptosis by Mediating Intramitochondrial Transport of Phosphatidic Acid. Cell Metab 18, 287–295 (2013).

59. Aaltonen, M. J. et al. MICOS and phospholipid transfer by Ups2–Mdm35 organize membrane lipid synthesis in mitochondria. Journal of Cell Biology 213, 525–534 (2016).

60. Ikeda, A., Imai, Y. & Hattori, N. Neurodegeneration-associated mitochondrial proteins, CHCHD2 and CHCHD10–what distinguishes the two? Front Cell Dev Biol 10, (2022).

61. Morgenstern, M. et al. Quantitative high-confidence human mitochondrial proteome and its dynamics in cellular context. Cell Metab 33, 2464–2483.e18 (2021).

62. Liu, X. et al. CHCHD2 up-regulation in Huntington disease mediates a compensatory protective response against oxidative stress. Cell Death Dis 15, 126 (2024).

63. Lisowski, P. et al. Mutant huntingtin impairs neurodevelopment in human brain organoids through CHCHD2-mediated neurometabolic failure. Nat Commun 15, 7027 (2024).

64. Köhler, G. & Milstein, C. Continuous cultures of fused cells secreting antibody of predefined specificity. Nature 256, 495–497 (1975).

65. Jiang, S. et al. TEFM regulates both transcription elongation and RNA processing in mitochondria. EMBO Rep 20, (2019).

66. Wredenberg, A. et al. Increased mitochondrial mass in mitochondrial myopathy mice. Proceedings of the National Academy of Sciences 99, 15066–15071 (2002).

67. Jiang, S. et al. Inhibition of mammalian mtDNA transcription acts paradoxically to reverse diet-induced hepatosteatosis and obesity. Nat Metab (2024) doi:10.1038/s42255-024-01038-3.

68. Mantas, I. et al. Genetic deletion of GPR88 enhances the locomotor response to L-DOPA in experimental parkinsonism while counteracting the induction of dyskinesia. Neuropharmacology 162, 107829 (2020).

69. He, Y. et al. Prosaposin maintains lipid homeostasis in dopamine neurons and counteracts experimental parkinsonism in rodents. Nat Commun 14, 5804 (2023).

70. Yang, L. & Beal, M. F. Determination of Neurotransmitter Levels in Models of Parkinson’s Disease by HPLC-ECD. in 401–415 (2011). doi:10.1007/978-1-61779-328-8_27.

71. Hubbard, K. E. et al. Determination of dopamine, serotonin, and their metabolites in pediatric cerebrospinal fluid by isocratic high performance liquid chromatography coupled with electrochemical detection. Biomedical Chromatography 24, 626–631 (2010).

72. He, Y. et al. Prosaposin maintains lipid homeostasis in dopamine neurons and counteracts experimental parkinsonism in rodents. Nat Commun 14, 5804 (2023).

73. Kaya, I. et al. Spatial lipidomics reveals brain region-specific changes of sulfatides in an experimental MPTP Parkinson’s disease primate model. NPJ Parkinsons Dis 9, 118 (2023).

74. Allen Institute for Brain Science. Allen Mouse Brain Atlas [adult]. Available from mouse.brain-map.org. Allen Institute for Brain Science (2011)

75. Filograna, R. et al. Modulation of mtDNA copy number ameliorates the pathological consequences of a heteroplasmic mtDNA mutation in the mouse. Sci Adv 5, (2019).

76. Wibom, R., Hagenfeldt, L. & von Döbeln, U. Measurement of ATP production and respiratory chain enzyme activities in mitochondria isolated from small muscle biopsy samples. Anal Biochem 311, 139–151 (2002).

77. Cox, J. & Mann, M. MaxQuant enables high peptide identification rates, individualized p.p.b.-range mass accuracies and proteome-wide protein quantification. Nat Biotechnol 26, 1367–1372 (2008).

78. Coudert, E. et al. Annotation of biologically relevant ligands in UniProtKB using ChEBI. Bioinformatics 39, (2023).

79. Tyanova, S. et al. The Perseus computational platform for comprehensive analysis of (prote)omics data. Nat Methods 13, 731–740 (2016).

80. Willforss, J., Siino, V. & Levander, F. OmicLoupe: facilitating biological discovery by interactive exploration of multiple omic datasets and statistical comparisons. BMC Bioinformatics 22, 107 (2021).

81. Milenkovic, D. et al. Preserved respiratory chain capacity and physiology in mice with profoundly reduced levels of mitochondrial respirasomes. Cell Metab 35, 1799–1813.e7 (2023).

82. Nolte, H., MacVicar, T. D., Tellkamp, F. & Krüger, M. Instant Clue: A Software Suite for Interactive Data Visualization and Analysis. Sci Rep 8, 12648 (2018).

