## Supplementary figures and images for "The CHCHD2-CHCHD10 protein complex is modulated by mitochondrial dysfunction and alters lipid homeostasis in the mouse brain"

### supplementary figures

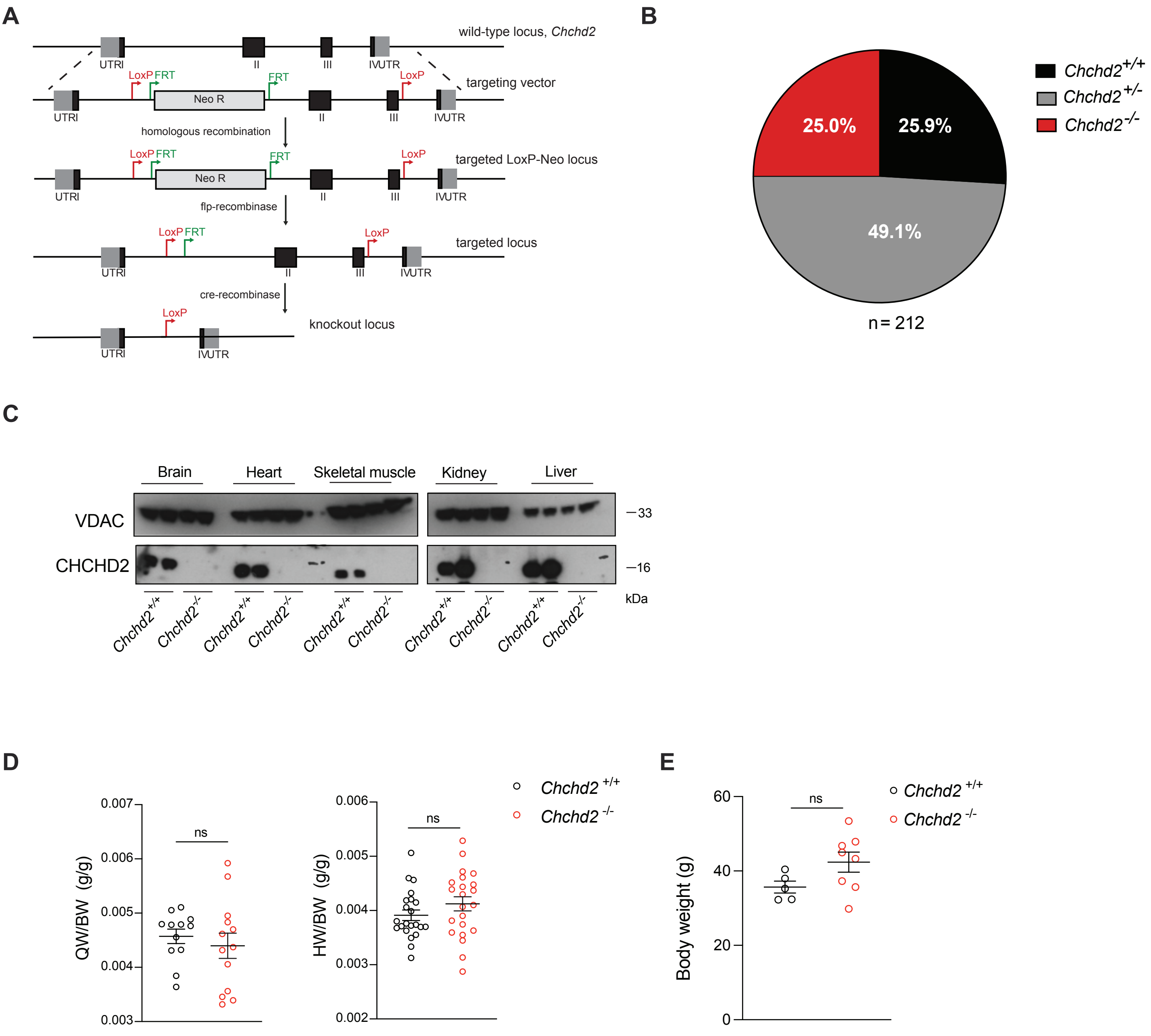

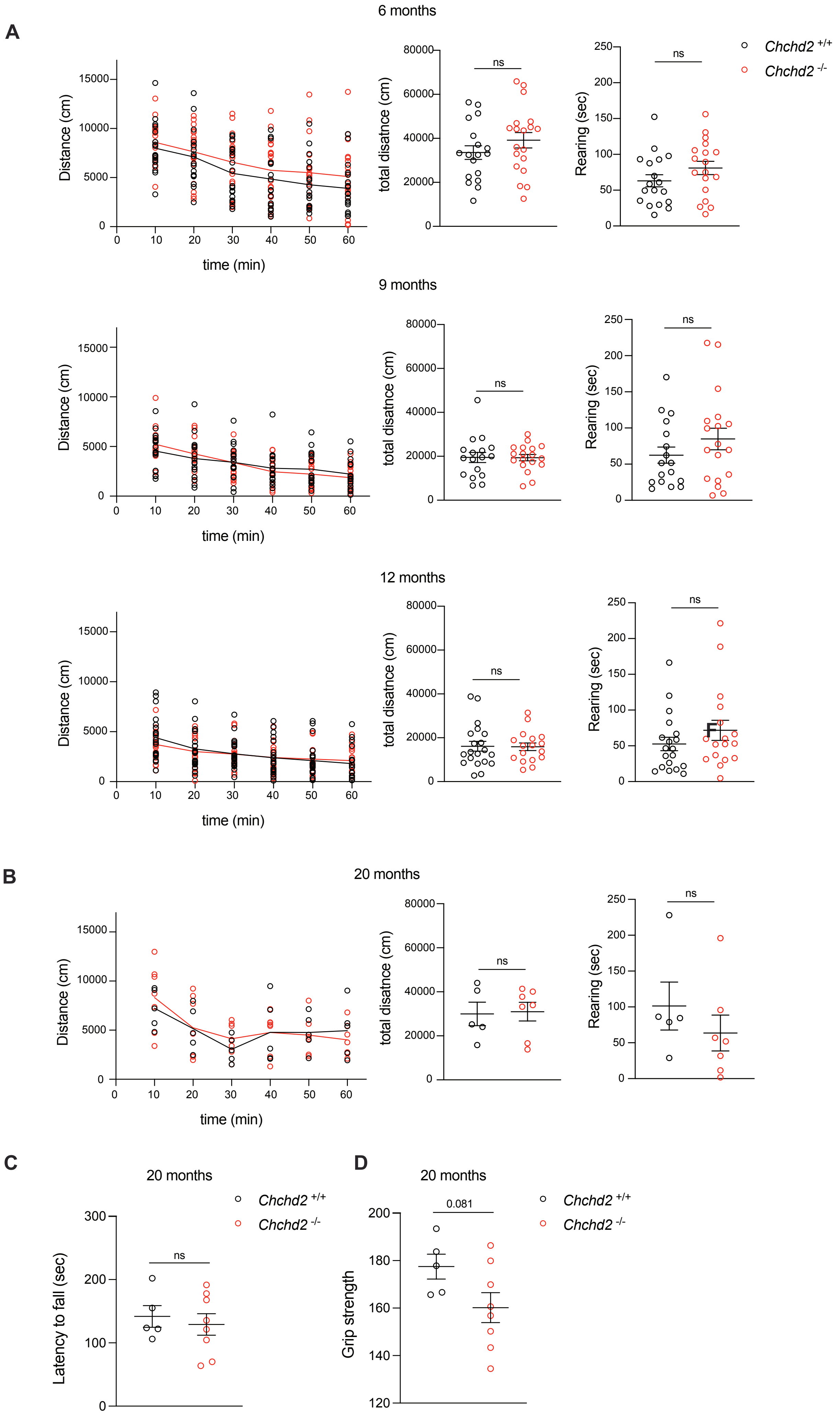

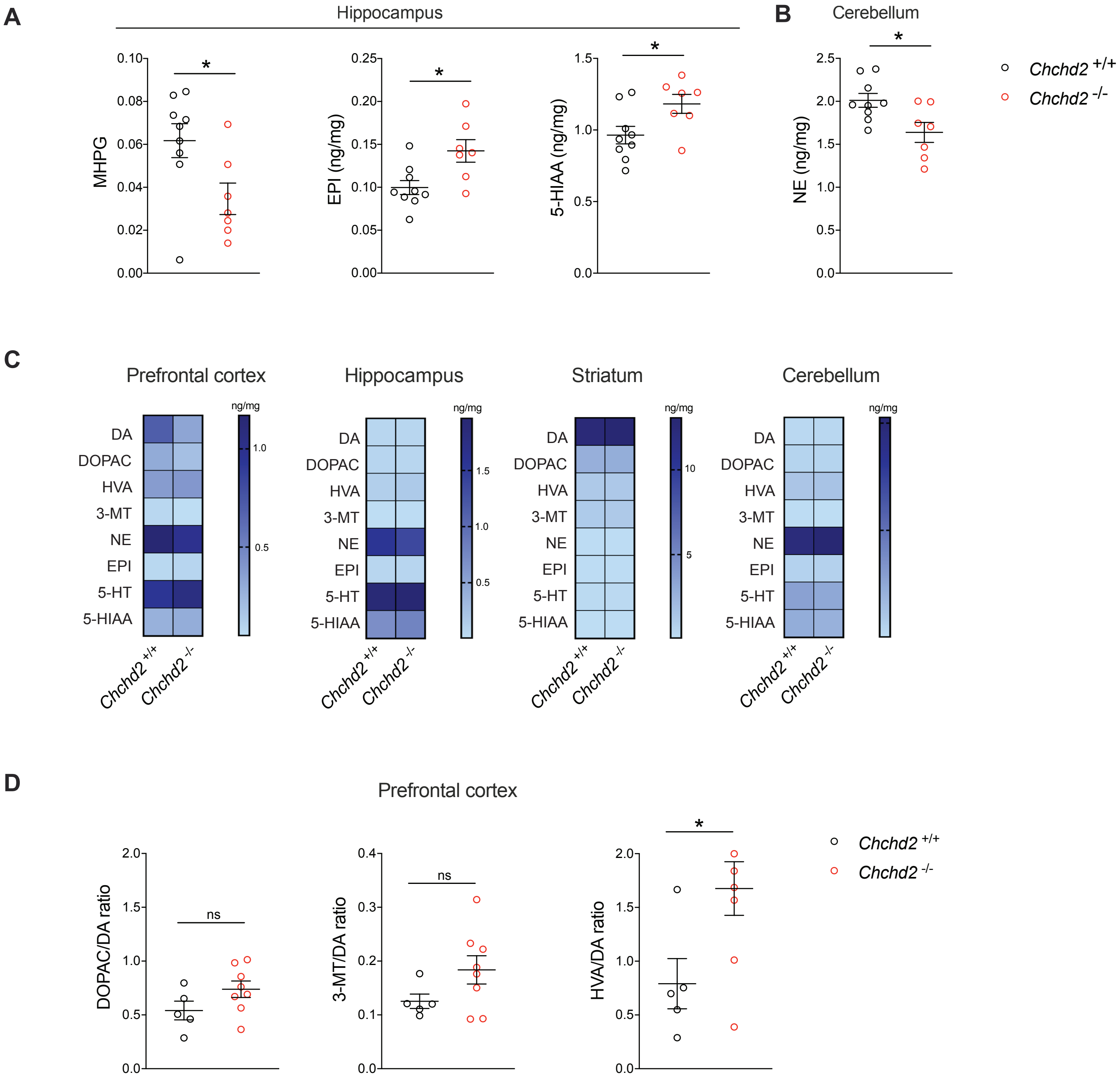

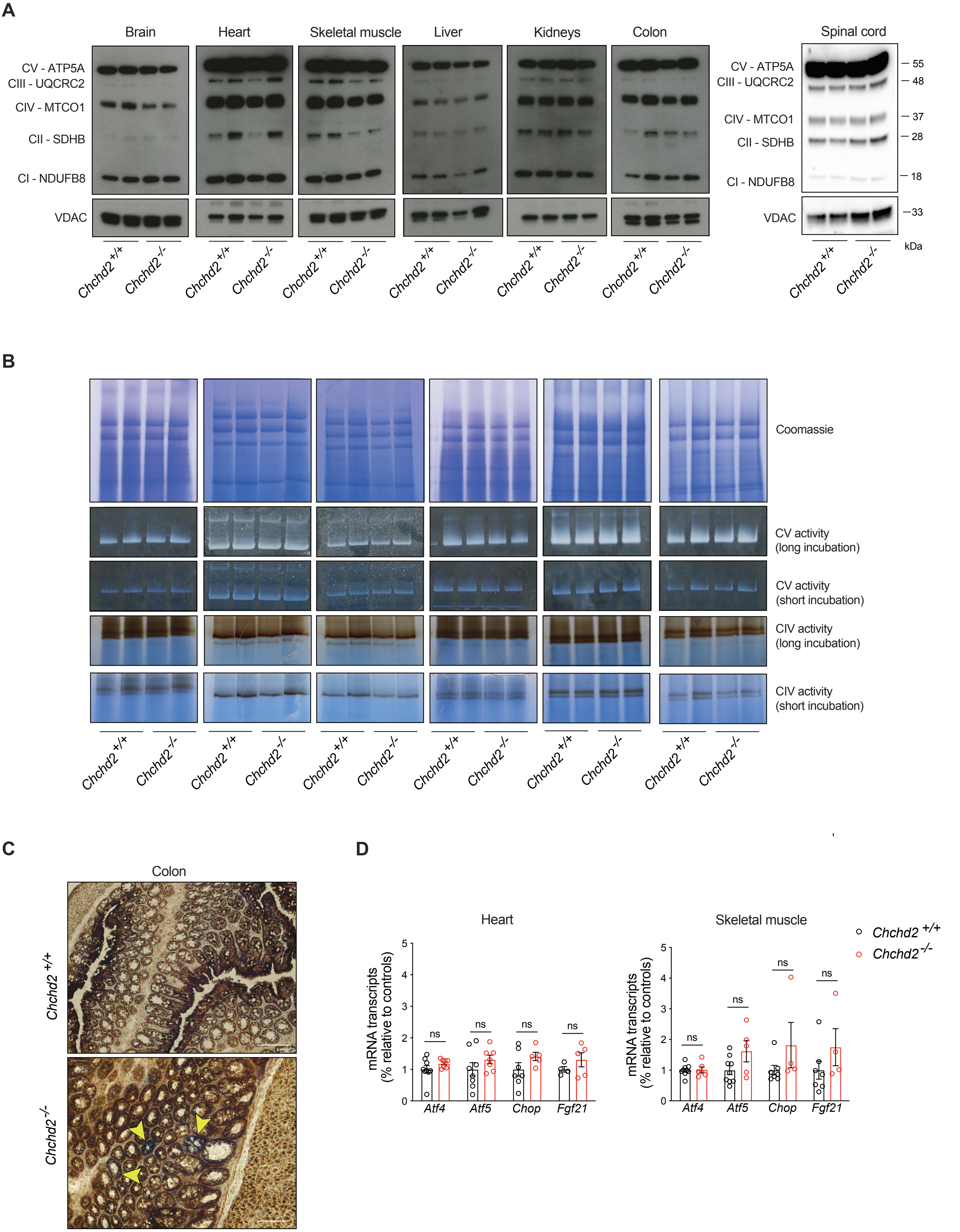

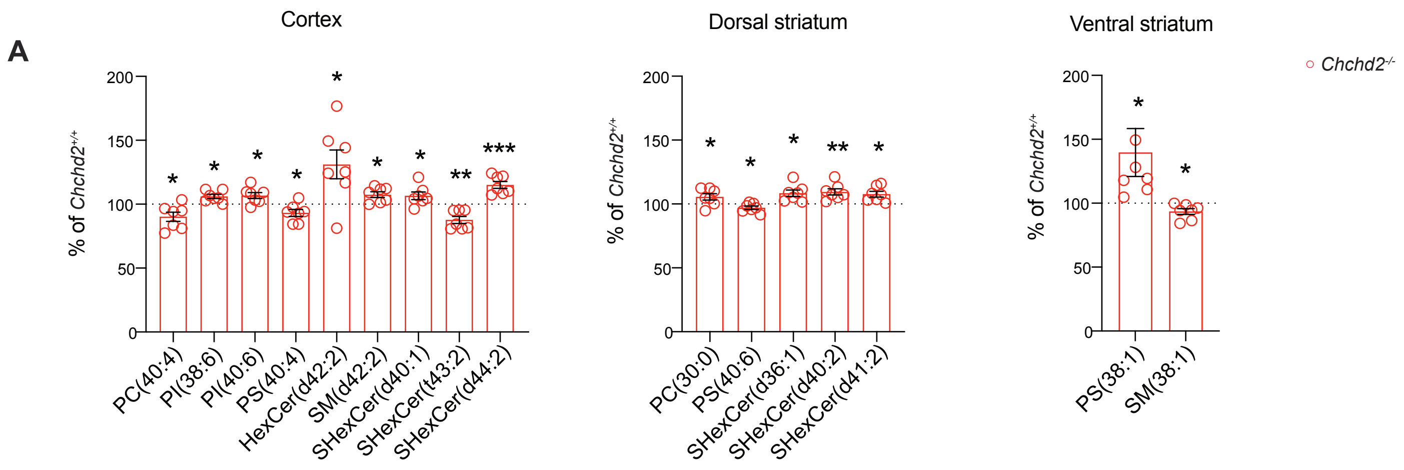

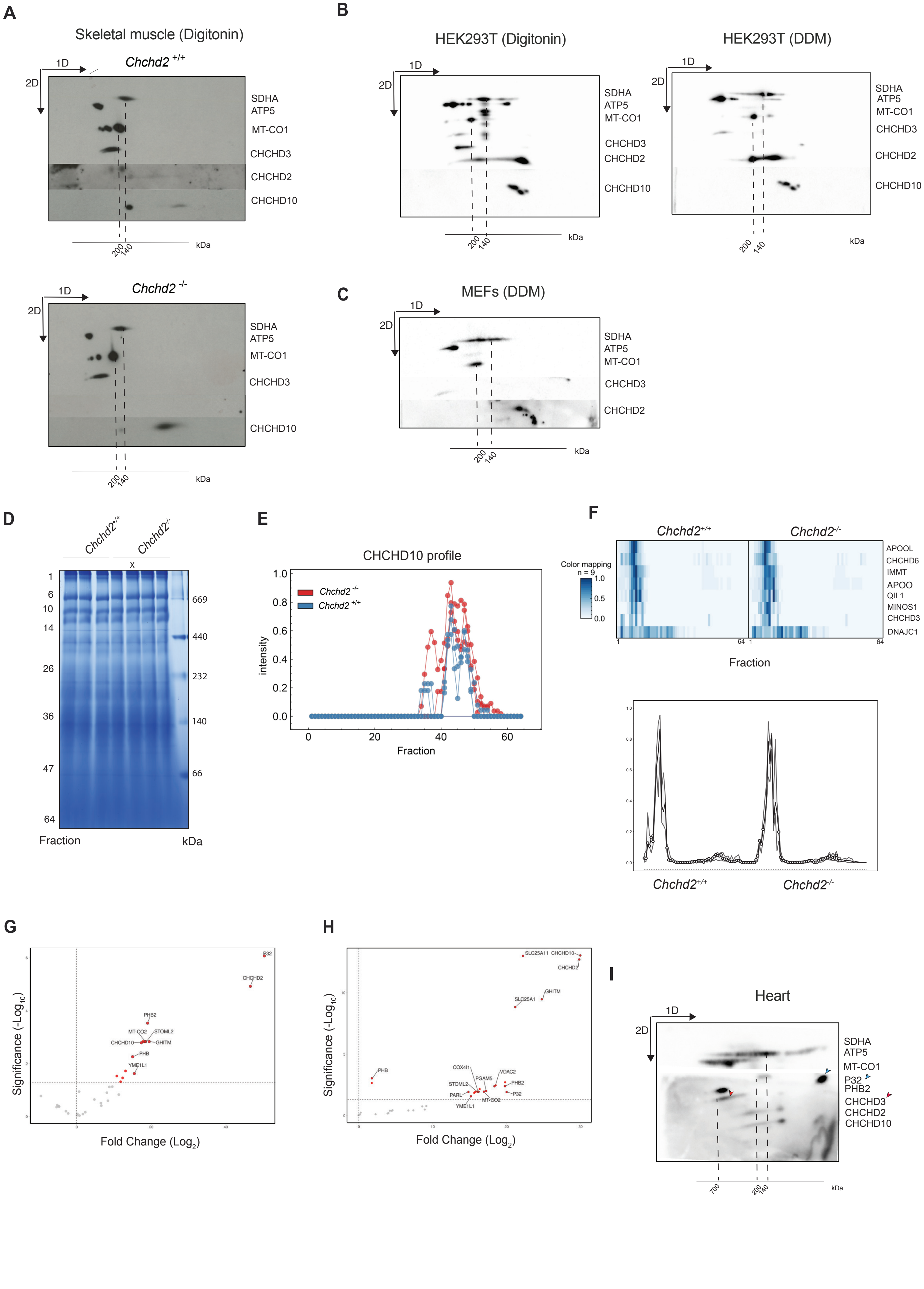

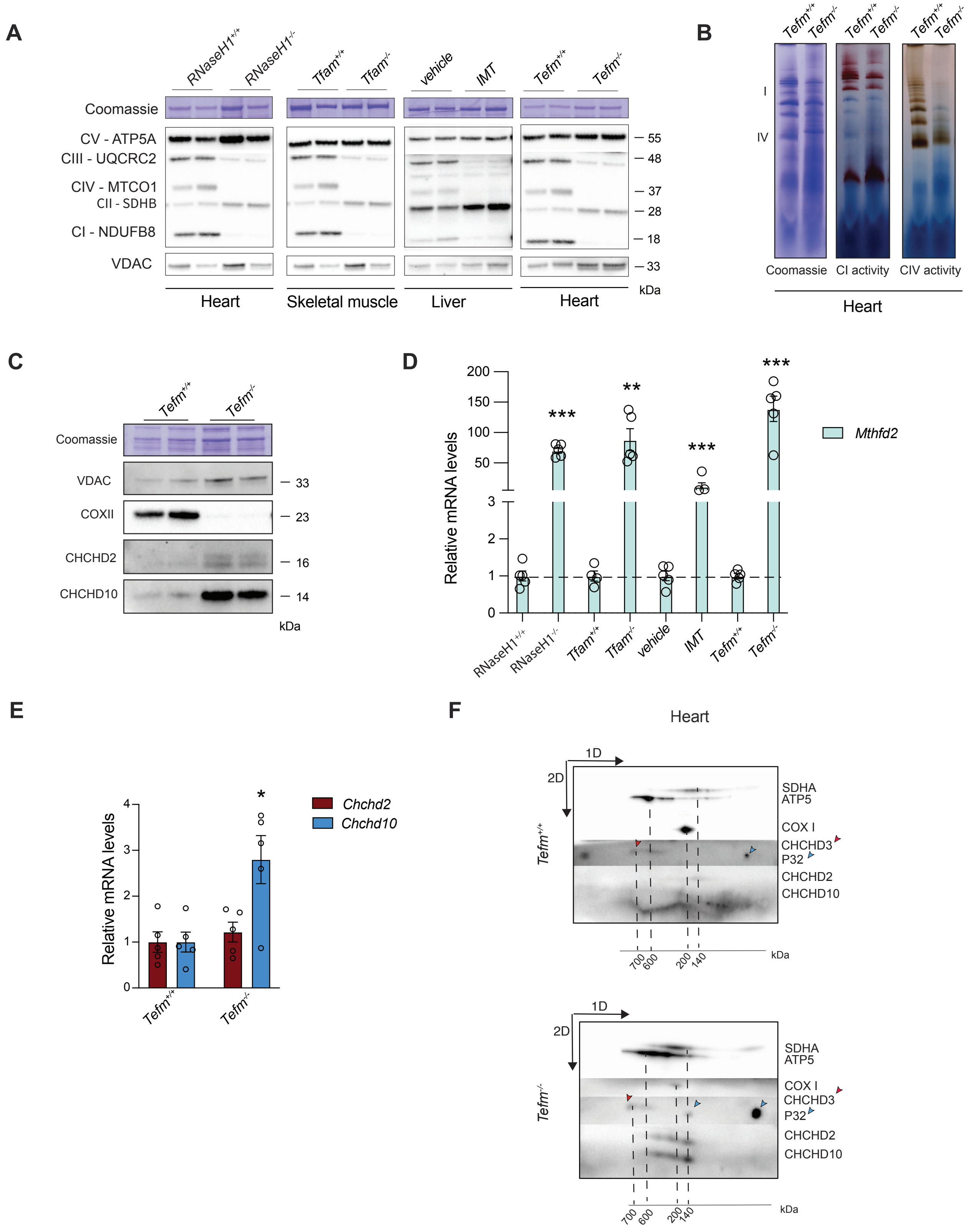

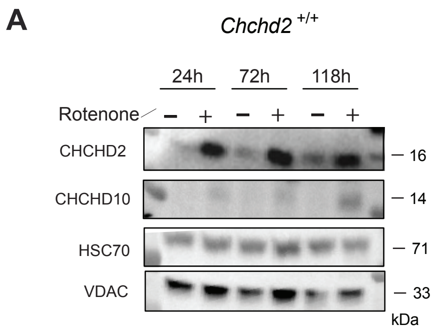
